# Energetic constraints shape the diversity of feasible ecological networks

**DOI:** 10.64898/2026.04.17.719283

**Authors:** Chengyi Long, Marco Tulio Angulo, C. Brandon Ogbunugafor, Ricard Solé, Serguei Saavedra

## Abstract

The relationship between energy supply and biodiversity is a longstanding question in ecology. Although a monotonic increase in diversity with energy availability is often assumed, unimodal species–energy relationships have been widely documented across ecosystems, and their origin from first principles remains unclear. Here, we develop a geometric framework that recasts ecological feasibility in explicitly energetic terms. By treating total energy supply as a system-level constraint on an energy-based network model, we define nested feasibility domains in the space of energy capture rates and quantify feasibility probabilities as their volume ratios. We show that the probability of initializing a feasible network increases monotonically and saturates with energy supply, whereas the probability of sustaining steady-state biomass follows a unimodal relationship—revealing a bounded energetic window within which network maturation is most likely. Extending this analysis to all candidate subcommunities via feasibility partitions, we find that different community sizes are most feasible at different energy levels, and that average diversity itself peaks at intermediate supply. Together, these results suggest that energetic constraints determine the diversity of ecological networks not through energy scarcity alone, but through the geometric interplay between external energy supply and internal energy exchange.

**Author Summary:** Why do many ecosystems show the highest biodiversity not where energy is most abundant, but at intermediate levels? This unimodal species–energy relationship has been documented across grasslands, wetlands, and rainforests, yet its origin from first principles has remained unclear. We approached this question by developing a simplified model that treats ecological networks as energy-processing systems. In this model, each species captures energy from the environment and exchanges it with others, and the total energy available to the network is explicitly limited. By measuring how the likelihood of species coexistence changes with energy supply within this framework, we found that while a minimum energy threshold is needed for any community to persist, too much energy can paradoxically reduce the chance of long-term coexistence. This creates a bounded energy window most favorable for community persistence. When we extended the analysis to all possible subsets of species, we found that different-sized communities are most likely to persist at different energy levels, and that overall expected diversity peaks at intermediate supply. These results suggest a possible geometric origin for why more energy does not always support more species, providing a theoretical baseline for connecting the structure of energy flow within networks to observed biodiversity patterns.

## Introduction

The success of life and the complexity it exhibits across scales are closely linked to its ability to harness free energy [1]. In ecosystems, species interactions and trophic structures form the ecological networks [2]. These networks represent the capture, exchange, and dissipation of energy [3, 4]. This has motivated the classical assumption that biodiversity increases monotonically with external energy supply, often identified as primary productivity. However, both the structure and the origin of this relationship remain the subject of ongoing controversy in the study of global biodiversity trends [5–8]. Across scales, the monotonic dependence between energy and diversity has been shown to break down: species-energy relationships often follow a unimodal curve, with a single hump observed in diverse ecosystems, including grasslands and wetlands, ponds, algal blooms, and rainforests [9–11]. What is the origin of these patterns? As noted in [5], they should be validated from first principles.

Following Boltzmann’s intuitions, Lotka proposed that natural selection should favor organisms that maximize energy flux [12]. This thermodynamic perspective has since inspired a broader view: that the organization of ecological networks is shaped not merely by external supply, but by the internal capacity to process and distribute energy [13–15]. In this view, the architecture of energy flow within a network—how energy is captured, routed, and dissipated across species—is more than a metaphor for trophic structure; it reflects genuine physical constraints on which community configurations can be sustained [3]. This raises a natural question: whether the structure of energy processing within ecological networks can help explain the origin of the diversity patterns described above—and, if so, how.

A related body of theoretical work has clarified important pieces of this problem, though each approach has left a distinct gap. Coexistence theory, grounded in resource competition and niche differentiation, has established that community persistence depends not only on external supply, but on how limiting resources are partitioned among species at equilibrium [16–20]—yet it does not treat total energy availability as an explicit, system-level constraint. Pioneering work in consumer-resource theory has shown that available energy flux and metabolic constraints shape community assembly and diversity [21, 22], but these analyses characterize system states along particular slices of resource space rather than offering a systematic treatment of the full space of configurations. Structural approaches have demonstrated that the geometry of species interactions defines feasibility domains—the regions of parameter space in which all species maintain positive abundances—in high-dimensional spaces [23–26]; however, the geometric methods developed are not directly applicable once a system-level energetic constraint is introduced. Together, these lines of work leave open a central question: how does a system-level energetic constraint shape the geometry of feasibility across the full space of possible ecological configurations?

To address this question, we develop a geometric framework that recasts ecological feasibility in explicitly energetic terms. Building on randomized model networks [27, 28], we reformulate the parameter space of ecological networks energetically and treat total energy supply as an explicit, system-level constraint. Within this energetic parameter space, we distinguish two developmental stages—initialization and maturation—that impose qualitatively different constraints on feasibility, and define two computationally tractable probabilities, ℙ_*I*_ and ℙ_*M*_, as volume ratios within the corresponding feasibility domains. These quantities allow us to systematically quantify how feasibility varies across network size, candidate community, energy exchange structure, and total energy supply, thereby revealing general energetic patterns across ecological networks. By grounding this analysis in a geometry-based formalism, our approach provides a foundation for future empirical investigations into the energetic boundaries of ecological persistence.

## Methods

### Energetic model of ecological networks

We model an ecological network of *S* populations as an open thermodynamic system, where biomass dynamics are governed by the balance of energy capture and dissipation and exchange, and all these energy fluxes are assumed to have per-capita forms (Fig. 1D-E). The rate of energy accumulation for population *i* is described by the following equation:

**Figure 1:**
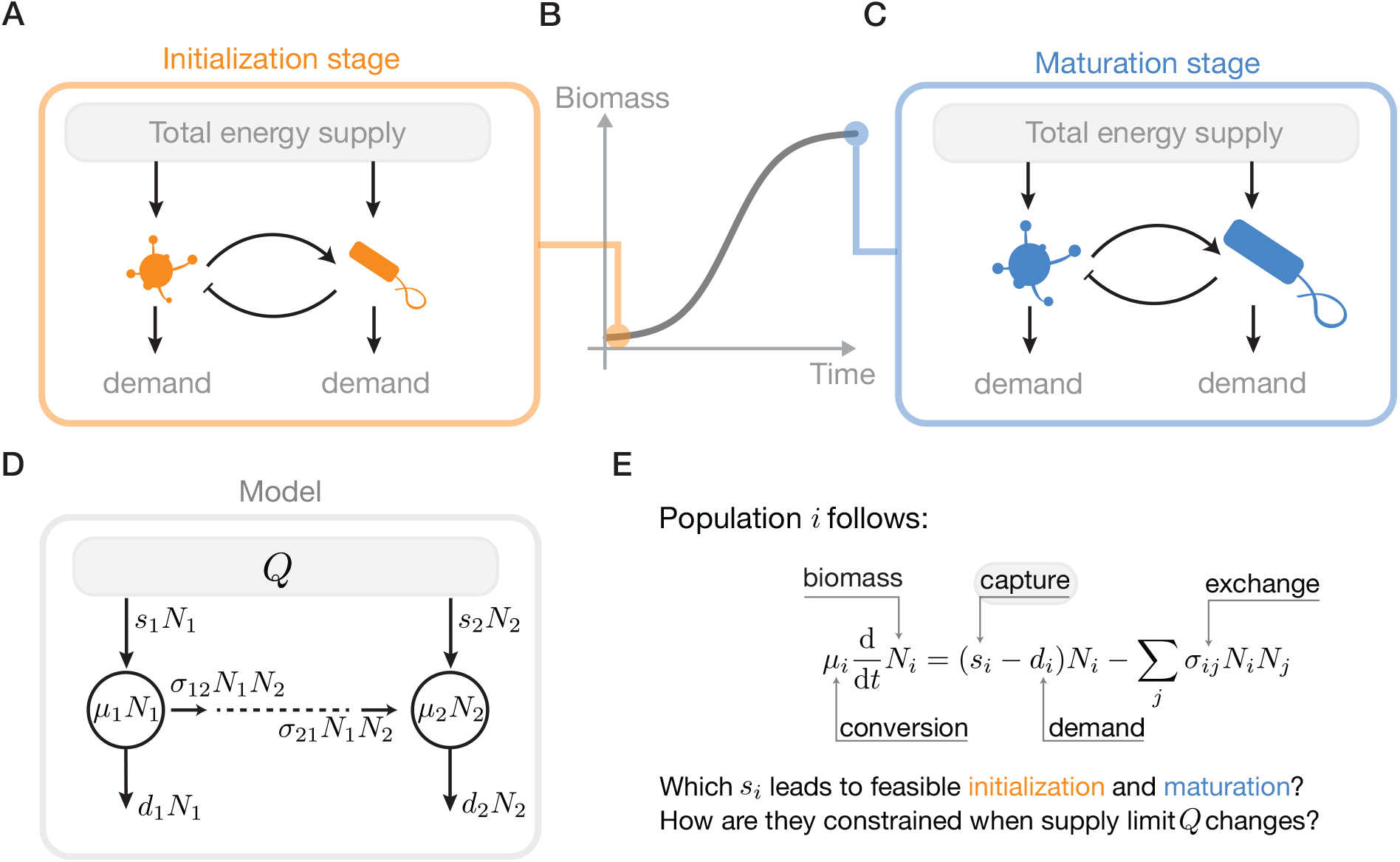
The energy flows and development of an ecological network. **A-C** Free energy is supplied from the external environment, captured by each population for their energy demands, and exchanged among them. As a result, they enable the biomass accumulation process of the network (**B**), particularly, the initialization (**A**) and maturation stage (**C**). **D** We model these energy flows as the product of corresponding mass-specific rates and biomass. The net energy flow into a population increases its biomass, which leads to the energy-based Lotka-Volterra dynamics (**E**). Notably, the total energy supply *Q* constrains the network’s energy capture.

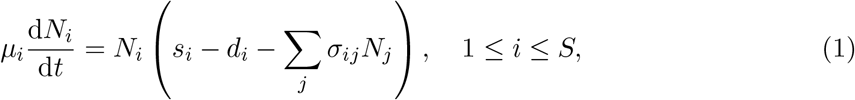

where *N*_*i*_ represents biomass ([mass]) and *µ*_*i*_ denotes energy density ([energy]·[mass]^−1^). The term *s*_*i*_ ≥ 0 denotes the mass-specific energy captured from the environment, while *d*_*i*_ ≥ 0 represents the energetic demands for maintenance ([power]·[mass]^−1^). The energy exchange matrix ***σ*** = (*σ*_*ij*_) quantifies the density-dependent cross-fluxes within the network ([power]·[mass]^−2^). Specifically, the diagonal *σ*_*ii*_ *>* 0 represents self-limitation and intrinsic losses (e.g., crowding effects), whereas off-diagonal elements *σ*_*ij*_ governs inter-specific energy exchanges, with positive and negative values indicating energy removal from or addition to the *i*-th population, respectively.

Therefore, the net energy balance is determined by: (1) *s*_*i*_*N*_*i*_, the total power captured from environmental resource gradients; (2) *d*_*i*_*N*_*i*_, the metabolic requirements for basal maintenance; and (3) ∑_*j*_ *σ*_*ij*_*N*_*i*_*N*_*j*_, the energy exchange between populations. It is noteworthy that, after rescaling by the energy densities *µ*_*i*_, Eq. (1) takes the form of a generalized Lotka–Volterra system, with intrinsic growth vector ***r*** ∝ (***s*** − ***d***) and interaction coefficients proportional to ***σ***. This connects our analysis directly to the feasibility literature, whose central question is which growth-rate vectors are compatible with coexistence for a given interaction matrix [23–26]. We retain this feasibility backbone, but give the parameters a more explicit energetic interpretation, which in turn allows us to impose concrete system-level energetic constraints in addition to the traditional positive-biomass condition.

### Feasibility domains for full communities

The emergence and persistence of an ecological network is determined by the alignment between the external energy availability and the network’s internal energetic structures. Given the steady-state biomass implied by Eq. (1), we therefore consider a maximal capacity of the environment to supply energy, defined as the *total energy supply rate Q* ([power]). Combined with internal structures, this boundary condition identifies four hierarchical conditions that relate mass-specific energy fluxes to the biological feasibility and energetic viability of the network (Fig. 2A):

**Figure 2:**
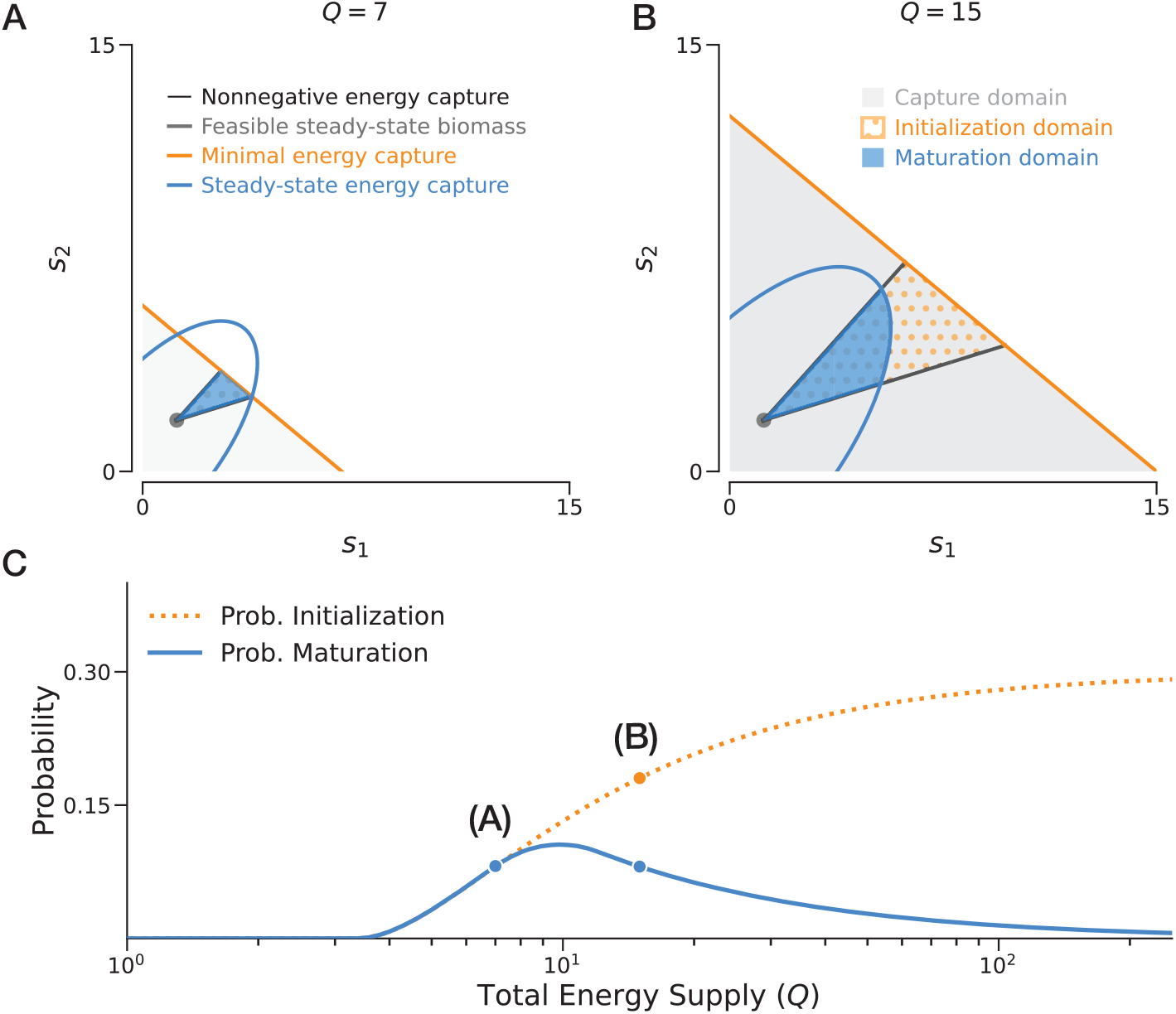
Constraints, feasibility, and probability in an example 2-population network. The network is parameterized with energy demand ***d*** = (1.20, 1.80)^⊤^, energy exchange matrix 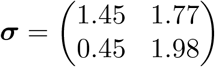, and minimal biomass vector ***N*** ^0^ = (1.00, 1.20)^⊤^. **A** In the parameter 0.45 1.98 space of ***s***, constraints related to energy fluxes (conditions i, ii, iii, iv) are represented by the linear and quadratic boundaries. **B** In the same space, we define the *capture domain* (gray shaded region), *initialization domain* (orange dots), and *maturation domain* (blue shaded region) as geometric intersections of the corresponding constraints. **C** Assuming ***s*** is uniformly distributed within the capture domain, probabilities are defined as volume ratios between the feasibility domains and the total possibility domain. As total energy supply *Q* varies, these probabilities may increase or decrease. Specifically, comparing *Q*_**A**_ = 7.0 and *Q*_**B**_ = 15.0, an increase in *Q* leads to a higher probability of initialization but a lower probability of maturation.

i. *Nonnegative energy capture:* Each population must receive a nonnegative rate of energy from the environment, i.e., *s*_*i*_ ≥ 0.
ii. *Feasible steady-state biomass:* At the steady state, each population must maintain a positive biomass: 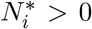 for all *i* [29, 30]. More generally, 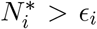, where *ϵ*_*i*_ *>* 0 is a positive threshold.
iii. *Minimal energy capture:* At the initialization stage, with a minimal biomass 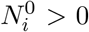 for each population, the total energy capture must not exceed the supply: 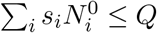.
iv. *Steady-state energy capture:* At the steady state, the total energy capture must remain within the environmental supply limit: 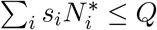.

For the case where all *S* populations coexist at equilibrium, these conditions define three nested domains within the space of energy capture rates 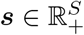 (Fig. 2B):

i. *The capture domain D*_*C*_(*Q*, ***N*** ^0^). This domain reflects the possible ways a total energy supply *Q* can be distributed without imposing any network-specific feasibility requirement. It is defined by conditions (i) and (iii): *D*_*C*_ = {***s*** ∈ ℝ^*S*^ | *s*_*i*_ ≥ 0 and 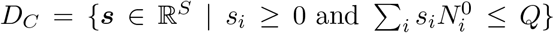}. Geometrically, this forms a simplex in the nonnegative orthant.
ii. *The initialization domain D*_*I*_(***σ, d*** | *Q*, ***N*** ^0^). This subset incorporates internal network structure by additionally requiring full coexistence at equilibrium. It is defined by the intersection of *D*_*C*_ and condition (ii): *D*_*I*_ = {***s*** ∈ *D*_*C*_ | ***N*** ^∗^ *>* 0}. This domain forms a convex polytope.
iii. *The maturation domain D*_*M*_ (***σ, d*** | *Q*, ***N*** ^0^). This further constrained subset requires that the steady-state energy capture remains feasible under the supply limit *Q*. It is defined by the intersection of *D*_*I*_ and condition (iv): 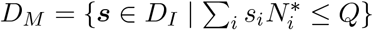. The condition 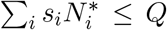 imposes a quadratic constraint. The nested structure *D*_*M*_ ⊆ *D*_*I*_ ⊆ *D*_*C*_ defines the feasibility landscape for ecological development under energetic constraints.

### Feasibility partitions for partial communities

To extend this framework beyond full coexistence, especially to cases of partial coexistence, we consider any candidate community 𝒞 ⊆ {1, …, *S*}. This requires generalizing the ***N*** ^∗^ = ***σ***^−1^(***s*** − ***d***) relationship to a boundary equilibrium supported on 𝒞. In Section 8 of S1 Text, we show that a similar construction can be rewritten for each 𝒞 by solving the complementarity condition. Under such generalization, condition (ii) requires positive biomass only for members of 𝒞, and condition (iv) evaluates the steady-state energy capture at the boundary equilibrium biomass.

In particular, the generalized version of condition (ii) can be written directly as a subset of ***s***-space [31]:

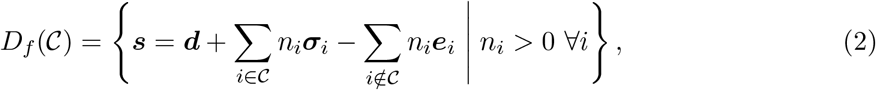

where ***σ***_*i*_ is the *i*-th column of ***σ*** and ***e***_*i*_ is the *i*-th unit vector. This representation allows us to naturally extend the definition of feasibility domains for partial coexistence. Specifically, for candidate community 𝒞, the initialization domain is

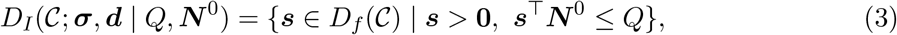

whereas the maturation domain is

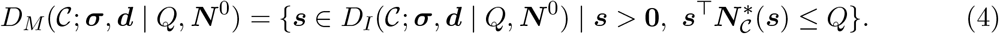

Here 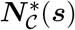 denotes the boundary equilibrium supported on 𝒞.

Across all 2^*S*^ candidate communities, the corresponding sets {*D*_*f*_ (𝒞)} induce a conic polyhedral partition of the ***s***-space around the full-coexistence region, with each direction of energetic input assigned to one candidate equilibrium support. For this reason, we refer to the resulting collection in (3) and (4) as *feasibility partitions*. The explicit derivations and proofs are provided in Section 8 of S1 Text.

Throughout this study, we assume that ***σ*** is Volterra dissipative [32], a standard restriction in Lotka–Volterra theory that implies global stability of the associated dynamics. Under this condition, ***s*** ∈ *D*_*f*_ (𝒞) is equivalent to the positive, stable coexistence of 𝒞. This reduces the analysis of coexistence to the analysis of feasibility.

### Probabilistic formulation and computation

Following the established framework of mapping ecological feasibility to geometric volumes [33], we treat the energy capture rates ***s*** as random variables. Given the inherent randomness of environmental inputs, we assume ***s*** is uniformly distributed over the capture domain *D*_*C*_—the *S*-dimensional simplex representing all physically possible resource-uptake configurations under a total supply *Q*. Under this assumption, the probabilities of initialization (ℙ_*I*_) and maturation (ℙ_*M*_ ) for full coexistence are defined as the relative volumes of their respective feasibility domains (Fig. 2C):

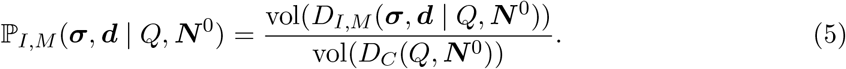

Similarly, we define the probability associated with any candidate community 𝒞 by replacing *D*_*I,M*_ with the corresponding initialization or maturation partition, *D*_*I,M*_ (𝒞; ***σ, d*** | *Q*, ***N*** ^0^).

The volume of *D*_*C*_ is determined analytically as 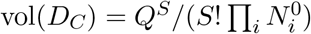. For the initialization domain *D*_*I*_, which is a convex polytope defined by linear inequalities, we compute the volume exactly using V-representation and H-representation decomposition algorithms [34].

For the maturation domain *D*_*M*_, the steady-state energy capture condition generally introduces a quadratic geometry described by the energy exchange matrix ***σ***. However, since we have assumed ***σ*** to be Volterra dissipative, it can be shown that *D*_*M*_ remains a convex body, allowing efficient volume estimation (Section 1 of S1 Text). To estimate vol(*D*_*M*_ ), we standardize the domain into a unified mathematical class (*InterPolyQuads*) by applying a linear transformation derived from the Cholesky decomposition of the symmetrized inverse of ***σ***. We then employ a Hit-and-Run Markov Chain Monte Carlo (MCMC) sampler [35] to generate uniform samples, using the Chebyshev ball (the largest inscribed ball) as stable starting points for the random walks. The final volume is estimated via a Multiphase Monte Carlo method [36] using 10^5^–10^6^ samples per domain. This computational pipeline was validated against analytical solutions for high-dimensional spheres, maintaining a relative error *<* 3% for dimensions as high as *S* = 12 (Section 2 of S1 Text).

### Model networks and simulation

To investigate general patterns of probabilities (ℙ_*I,M*_ ) as a function of total energy supply (*Q*), we broadly follow the random-matrix approach widely used in community ecology [27, 37, 38]. However, rather than directly drawing from certain distributions, we further constrain the sampled (***σ, d, N*** ^0^) to match the assumptions in our framework. Specifically, we require the energy transfer efficiency to be at most 1 for opposite-sign *σ*_*ij*_, *σ*_*ji*_ pairs, we require ***σ*** to be dissipative, meaning that (***σ*** + ***σ***^⊤^)*/*2 is positive definite [32], and we require both ***d*** and ***N*** ^0^ to remain strictly positive. Together, these restrictions define the ensemble of model networks analyzed in this study.

To generate such matrices, we use a four-step procedure: (1) *Sampling* : elements *σ*_*ij*_ are first drawn from Normal(0, 1); (2) *Efficiency* : for any opposite-sign pair violating the efficiency constraint, the corresponding absolute values are swapped so that |*σ*_*ij*_| ≤ |*σ*_*ji*_| whenever *σ*_*ij*_ *<* 0 and *σ*_*ji*_ *>* 0; (3) *Dissipativity* : diagonal elements are shifted, *σ*_*ii*_ ← *σ*_*ii*_ + *c*, so that the minimum eigenvalue of the symmetric part is *σ*_0_ *>* 0; and (4) *Scaling* : the full matrix is multiplied by a factor *s*_*σ*_. Minimal biomass ***N*** ^0^ and energy demands ***d*** are then drawn independently from normal distributions centered at *N*_0_ and *d*_0_, respectively, and are ensured to be positive.

For the analyses reported here, we generated 100 independent random networks with *S* = 8 populations, utilizing parameter values: *s*_*σ*_ = 1.0, *σ*_0_ = 0.5, *N*_0_ = 1.0, and *d*_0_ = 1.0. To evaluate sensitivity to network size, we down-sampled these networks to *S* ∈ {2, 4, 6}. In Section 4 of S1 Text, we further tested the sensitivity of the results by rescaling ***σ, d***, and ***N*** ^0^ by factors of 2 and 0.5, by varying the diagonal-shift parameter *σ*_0_, and by varying the network connectance. These changes affect the probabilities quantitatively, but preserve the same qualitative patterns associated with our results.

### Functional data analysis

To extract general patterns between feasibility probability and total energy supply across randomized networks, we treat each realization as a probability curve ℙ(*Q*; ***θ***), where ***θ*** = {***σ, d, N*** ^0^} denotes the ecological parameters of that network. Therefore, we deploy functional data analysis (FDA) to estimate the general trend and variation from these probability curves.

When calculating each realization (***θ***), the probability curve was first evaluated on a ***θ***-dependent grid of *Q* values, which automatically covers qualitative patterns of the curve (Section 5.3 of S1 Text). When analyzing these curves, we first linearly interpolated each curve onto a common logarithmically spaced grid of *Q*.

We then clamp probabilities to the interval [*ϵ*, 1−*ϵ*] with *ϵ* = 10^−3^ and apply the logit transform

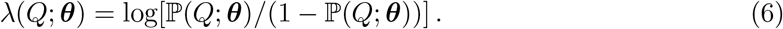

For each value of *Q*, we compute the pointwise mean *µ*_*λ*_(*Q*) and standard deviation *σ*_*λ*_(*Q*) across the ensemble in this transformed space. The aggregated trend is then mapped back to probability space using the inverse logit. Specifically, we report the the inverse-logit of the pointwise mean in logit space as

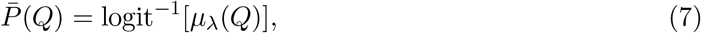

together with a one-standard-deviation band in logit space defined by

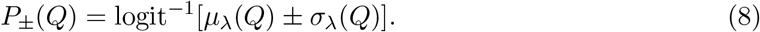

This transformation keeps both the central trend and uncertainty bands within the biologically meaningful range [0, 1] while allowing asymmetric uncertainty near the boundaries. Similarly, diversity curves (⟨*D*⟩) were summarized directly on their original scale by taking the pointwise arithmetic mean across interpolated *Q* grid.

To verify that the ensemble-level patterns also hold at the level of individual networks, we further quantified the degree to which each ℙ_*I*_(*Q*) curve followed a saturating pattern and each ℙ_*M*_ (*Q*) curve followed a unimodal pattern using summary scores defined in Section 3 of S1 Text.

## Results

### Energetic regimes in a two-population network

To illustrate the geometry of feasibility domains under varying total energy supply, we first analyze an example two-population network (*S* = 2) whose parameters are randomly sampled. As the total energy supply *Q* increases, three distinct phases characterize the network’s feasibility (Fig. 2C). In the low-energy regime (*Q <* 3.4), both initialization (ℙ_*I*_) and maturation (ℙ_*M*_ ) remain zero, defining a critical threshold *Q*_*c*_ = 3.4, below which insufficient energy supply leads to infeasible network. In the intermediate-energy regime (3.4 *< Q <* 10.0), both probabilities rise with increasing supply. However, a divergence occurs in the high-energy regime (*Q >* 10.0): while ℙ_*I*_ continues to increase and eventually saturates, ℙ_*M*_ exhibits a distinct decay after reaching a peak at *Q*_opt_ = 10.0. This *unimodal response* of ℙ_*M*_ suggests that excessive energy supply can paradoxically reduce the likelihood of long-term network maturation.

The geometrical interplay between energy capture constraints and network structures explains these energetic regimes (Fig. 2A-B). The initialization probability ℙ_*I*_ is constrained solely by the intersection of the feasible steady-state biomass (a fixed cone defined by ***N*** ^∗^ *>* **0**) and the minimal energy capture constraint 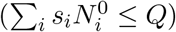. As total energy supply increases, the critical threshold *Q*_*c*_ is the point where minimal energy capture first overlaps with the feasible steadystate region. In general, Section 5 of S1 Text shows that *Q*_*c*_ ≤ ***d***^⊤^***N*** ^0^, with equality holding in many cases. As *Q* continues to grow, the relative volume of this intersection (i.e. initialization domain) approaches a geometric limit determined by the angular spread of the steady-state cone, leading to the observed saturation of ℙ_*I*_. This saturation behavior is consistent with the concept of structural stability [23].

In contrast, the probability of maturation (ℙ_*M*_ ) is additionally shaped by the steady-state energy capture constraint 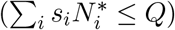. Because steady-state biomass ***N*** ^∗^ typically increases in response of larger *Q* (thus larger ***s***), this constraint becomes more stringent and grows more slowly as *Q* increases, but is only relevant when *Q* is sufficiently large. Consequently, ℙ_*M*_ undergoes a transition: at lower *Q*, the maturation domain is bounded by the minimal energy capture constraint (Fig. 2A), but as *Q* increases, the steady-state energy capture constraint becomes dominant, progressively prunes the feasible region (Fig 2B). This transition creates the optimal energy zone around *Q*_opt_, at which the network maturation is most feasible. Detailed analytical derivations of *Q*_*c*_ and *Q*_opt_ are provided in Section 5 of S1 Text.

### Feasibility patterns and drivers for multi-species full communities

To test the generality of these energetic regimes, we analyzed ensembles of sampled model networks with sizes *S* ∈ {2, 4, 6, 8}. For each network size, we considered 100 independent realizations and focused on the feasibility of the full community (i.e. all species coexist). We extracted robust general trends from these ensembles using functional data analysis (FDA). The fundamental relationships identified in the two-population case – a monotonic saturation for initialization (ℙ_*I*_) and a unimodal response for maturation (ℙ_*M*_ ) – persist as robust features across these random networks with diverse system parameters (***σ, d, N*** ^0^). Among all network sizes, initialization requires a critical total energy supply *Q*_*c*_, above which ℙ_*I*_ rises and eventually plateaus (Fig 3A). In addition, maturation remains constrained within a narrower window of energy supply, characterized by an optimal energy supply level *Q*_opt_, at which the probability of maturation reaches its peak. These patterns are consistent with individual networks (Section 3 of S1 Text), and are qualitatively robust to the sampling choices or hyperparameters to generate the ensemble (Section 4 of S1 Text).

**Figure 3:**
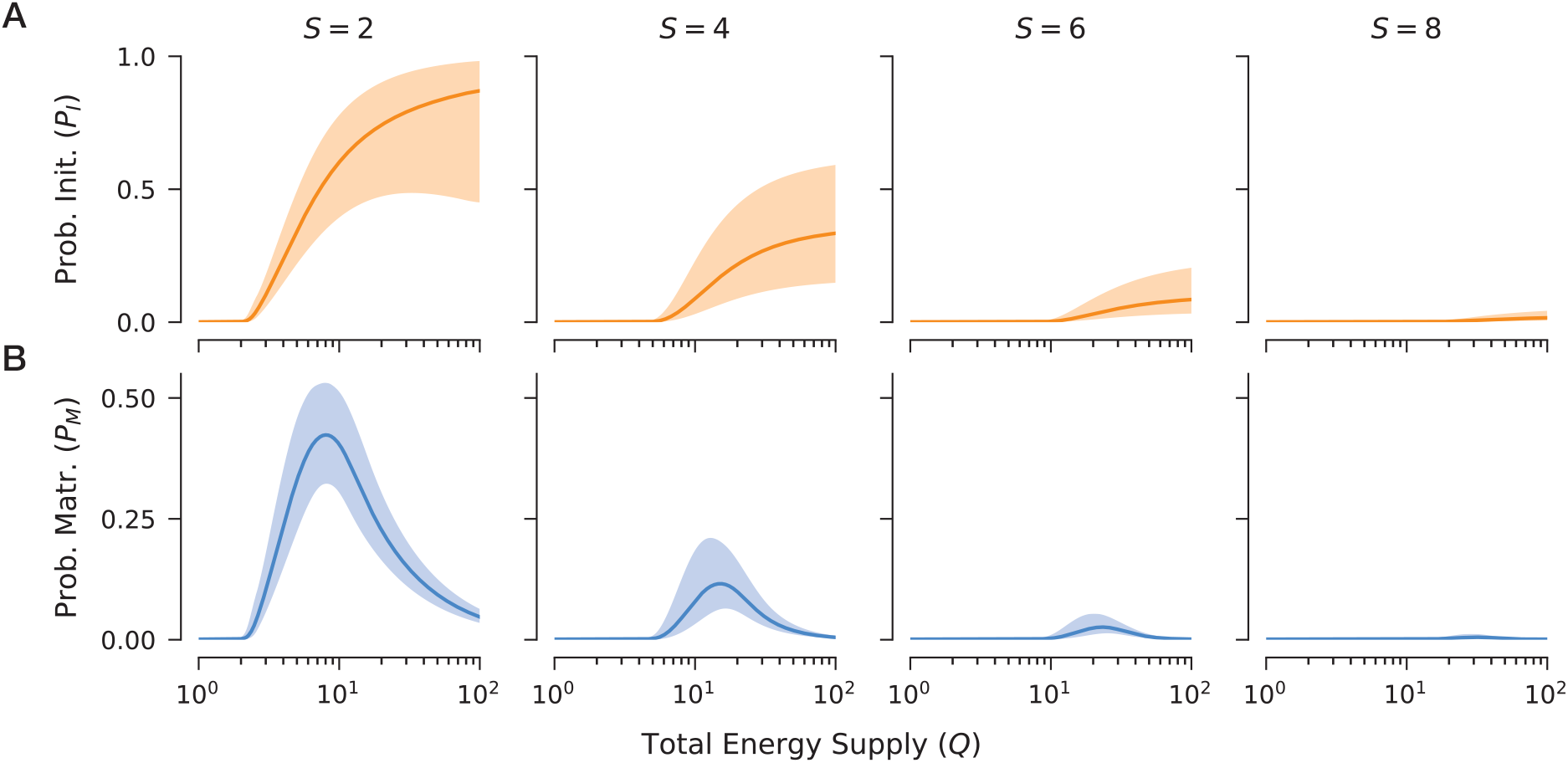
General patterns of feasibility probabilities under varying total energy supply and network sizes. We compute the expected probabilities of initialization and maturation for networks of varying total energy supply (*Q* ∈ [10^0^, 10^2^]) and network size (*S* ∈ {2, 4, 6, 8}), from randomly sampled ecological parameters (***σ, d, N*** ^0^). Each panel represents 100 independent replicates. **A** The probability of being feasible at the initial stage is a monotonically increasing function of energy supply, gradually saturating at higher *Q* values. **B** The probability of being feasible at the maturation stage exhibits a unimodal relationship with energy supply, peaking within an optimal energy range. In both cases, probabilities decrease as network size *S* increases, regardless of *Q*. For each replicate, probabilities are evaluated across 500 logarithmically spaced values of *Q*. Solid lines and shaded areas represent the pointwise mean and one-standard-deviation band in logit space, respectively, estimated via functional data analysis (see Methods). The saturation and unimodal patterns are robust at individual replicate level with high statistical scores (Section 3 of S1 Text). Although we focus here on a specific parameter setup, qualitative trends are preserved across a broader parameter space, as demonstrated by a comprehensive sensitivity analysis in Section 4 of S1 Text.

To ensure these patterns are not artifacts of the linear assumptions in our primary framework, we tested the robustness of our results by incorporating density-dependent capture rates to mimic saturating resource-uptake kinetics [16]. As a result, the relationship between steadystate biomass (***N*** ^∗^) and energy capture rates (***s***) shifts from a linear to a saturating nonlinear form. We find that while this self-regulatory feedback leads to a non-zero limiting probability of maturation at extremely high energy supply (*Q*), the characteristic unimodal response remains a robust feature within the low-to-intermediate *Q* regime. This confirms that the identified energetic window is a general property of energy-constrained networks, persisting independently of the functional linearity in the energy–biomass relationship (see Section 7 of S1 Text for detailed derivations and results).

The primary determinant of feasibility across the ensemble is network size. Both ℙ_*I*_ and ℙ_*M*_ decrease significantly as the number of populations *S* increases, regardless of the energy supply *Q* (Fig. 3). This decline suggests an inherent “energetic cost of complexity”, where longer internal chains of energy flux – associated with larger networks – face increasingly narrow paths to feasibility. This finding aligns with empirical observations regarding limited length of trophic chains in natural ecosystems [39]. However, our sensitivity analyses reveal that other parameters can modulate this relationship, allowing for a more nuanced quantification of feasibility (Section 4 of S1 Text). For example, increasing the minimal biomass (***N*** ^0^) shifts both *Q*_*c*_ and *Q*_opt_ to higher energy levels, while also increases the maximum maturation probability at *Q*_opt_. Similarly, increasing the magnitude of energy exchanges (***σ***) leaves *Q*_*c*_ and ℙ_*I*_ unaffected, but shifts *Q*_opt_ and ℙ_*M*_ (*Q*_opt_) upward. While these results are analyzed by isolating the impact of one parameter *ceteris paribus*, they could be combined to provide a mechanistic yardstick to evaluate how potentially countervailing factors could collectively determine the feasibility of multi-species ecological networks.

### Feasibility and average diversity across partial and full communities

To examine how the energetic patterns change once partial coexistence is allowed, we extended the analysis from full community to all candidate communities of a network. For a network of size *S*, there are 2^*S*^ candidate communities in total, and exactly 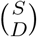 of them have community size *D* = |𝒞|. Focusing on *S* = 6, we summarized the maturation probabilities of all candidate communities by their size *D* ∈ {1, 2, 3, 4, 5, 6} (Fig. 4A). Similar results for *S* ∈ {2, 4, 8} are provided in Section 8 of S1 Text.

**Figure 4:**
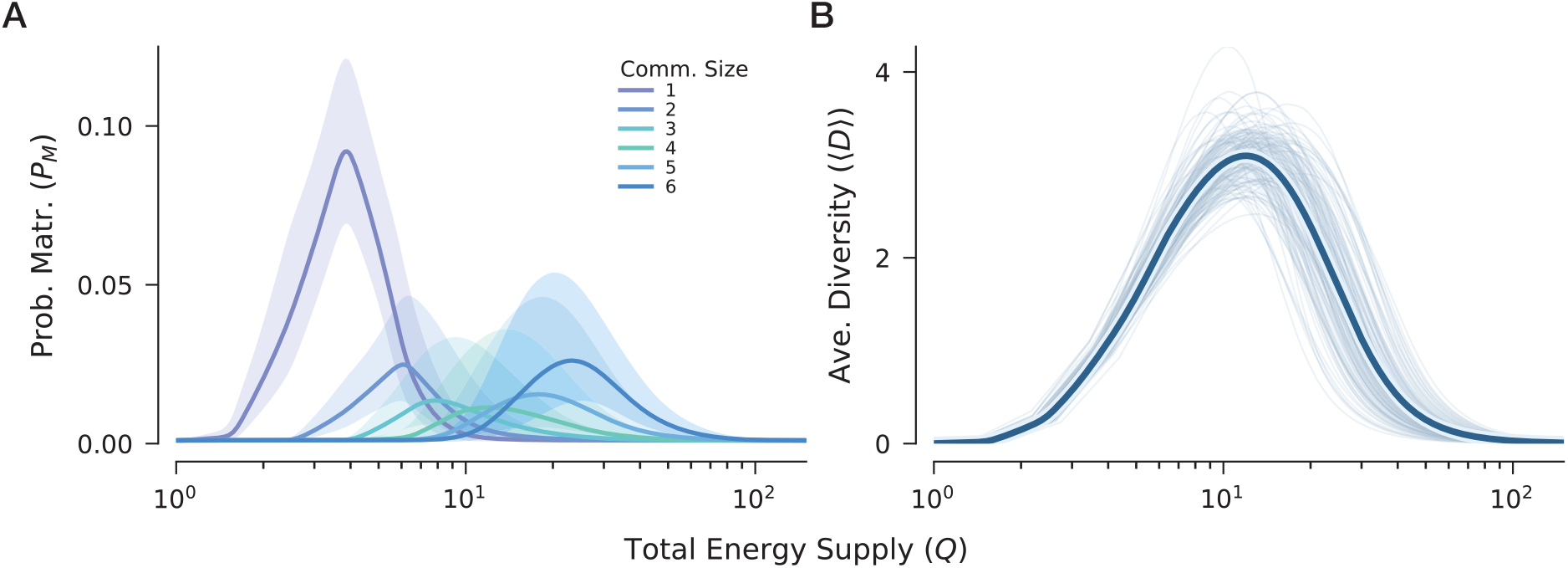
Probabilities of maturation and average diversity across full and partial coexistence. We extended the analysis from only full-coexistence to all candidate communities for model networks with size *S* = 6. For each network, all 2^6^ − 1 = 63 candidate communities were enumerated and grouped by community size 1 ≤ *D* ≤ 6, with 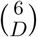 communities in each size class. For each candidate community, we computed the probability of maturation over 500 logarithmically spaced values of total energy supply *Q* ∈ [10^0^, 10^2^]. Panel **A** was summarized using the logit-based functional data analysis described in Methods, whereas panel **B** was summarized on the original richness scale after interpolation onto the same common *Q* grid. Each panel represents 100 independent network realizations. **A** Expected probability of maturation as a function of total energy supply for different community sizes. Across all sizes, probability of maturation retains a unimodal relationship with energy supply, but the corresponding energetic window shifts upwards as community size increases. **B** Average diversity, defined as the expected community size weighted by both the number of candidate communities 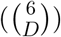 and their mean probabilities of maturation (*P*_*M*_ (*Q*)) at each *D*. Although medium-sized communities (*D* ∈ {3, 4}) have low mean probability of maturation, they are both combinatorial numerous and favored at intermediate energy windows. Due to this contribution, average diversity peaks at an intermediate *Q* lower than the *Q*_opt_ for full coexistence.

Across all community sizes, the probability of maturation retains the same qualitative unimodal relationship on total energy supply observed for full coexistence (Fig. 4A). However, the location of this energetic window shifts systematically with community size: smaller communities reach their highest maturation probability at lower supply levels, whereas larger communities require progressively higher *Q* to become most feasible. Thus, increasing energy supply does not elevate the feasibility of all candidate communities simultaneously; instead, it sequentially favors communities of different sizes across distinct energetic ranges on the logarithmic scale.

To translate these community-level patterns into a network-level quantity, we defined the average diversity as the expected community size, weighting each community size by both its number of candidate communities and their corresponding maturation probabilities. This aggregation also reveals a unimodal relationship with total energy supply (Fig. 4B). Importantly, the energy level that maximizes average diversity is lower than the optimal energy supply for full coexistence. This difference arises because intermediate-sized communities, although individually less likely to mature than the full community, are far more numerous. When these two factors are combined, the collective contribution of many moderately feasible, mid-sized communities dominates the network-level average diversity at intermediate energy supply. In this sense, the energy level that maximizes expected diversity is not the same as the one that maximizes the feasibility of the full-coexistence state.

## Discussion

Understanding the struggle for life as a struggle for free energy offers a unifying framework for investigating the emergence of complex living systems. It is plausible that these systems demonstrate emergent capacity to develop and persist under conditions of energy scarcity. This raises a fundamental and unresolved question: are there optimal energy levels at which life can thrive most effectively? In other words, can both insufficient and excessive energy availability reduce the likelihood of observing complex living systems? A low energy supply may impose a minimal threshold for persistence, reflecting the unavoidable dissipation costs inherent to all living systems. Conversely, an excessive energy supply may overwhelm internal regulatory structure, suggesting limits to the capacity of organisms or communities to manage internal energy exchange. Together, these considerations point to the possibility of a bounded “energy window” within which the system is most feasible.

To reveal this energy window quantitatively, we introduced an energy-based formalism that mapped ecological dynamics onto a geometrical representation. This representation was built on previous geometric treatments of feasibility domains [23, 26, 30], while recasting them in explicitly energetic terms. By leveraging advanced stochastic methods to estimate high-dimensional feasibility domains, our approach enabled a rigorous exploration of how the interplay between external input and internal exchange governed network feasibility. More broadly, this geometric formulation offered a useful framework for analyzing feasibility in community ecology and related complex systems, complementing previous sampling-based and geometric approaches [40, 41].

Following this modeling and computation framework, we first analyzed the feasibility of full communities across ensembles of model networks. Results showed that initialization was gated by a minimum energy requirement and eventually reached a feasibility plateau. In contrast, the unimodal response of maturation probability revealed an optimal regime of energy supply at which the full community was most likely to persist. This qualitative picture remained under a saturating, density-dependent uptake formulation, even though the probability of maturation approached a nonzero limit at high energy supply. Extended to all candidate communities, the same unimodal pattern remained for maturation probabilities, but their optimal total energy supply progressively increased with community size, indicating that communities of different sizes were most feasible at different energy levels. Collectively, the energy level that maximized average diversity of the system was lower than the one that maximized the feasibility of the full community. Overall, these findings suggested that external energy supply directly determined feasibility, whereas the corresponding energy window was shaped by the ecological network’s internal structure of energy transfer and utilization [30], resonating with broader views in community ecology [42, 43].

In addition to energy supply, network size and community size emerged as key determinants of feasibility, and therefore as mediators of the energy–diversity relationship revealed by our ensemble analysis. Within a given network, larger candidate communities generally required higher energy supply to become most feasible, whereas intermediate community sizes were combinatorially far more numerous (Fig. 4A). At the aggregate level, increasing *Q* expanded the set of admissible energy allocations and shifted which community supports could be realized, but did not translate monotonically into higher expected diversity. Instead, average diversity peaked at intermediate energy supply, where the collective contribution of many moderately feasible, intermediate-sized communities was greatest (Fig. 4B). In this sense, our results complement long-standing discussions of limited trophic length and food-web size [39]. Rather than treating these patterns as simple consequences of resource scarcity alone, our framework suggests that diversity limitations may also reflect structural constraints [on how energy can be distributed across feasible ecological networks]. Although this conclusion was derived from a simple model, it provides a concrete baseline for testing how these mechanisms scale in larger and more realistic ecological networks.

The Eqs. (2), (3), and (4) partition capture rate space (***s***-space) by feasible candidate community 𝒞. In particular, this geometry directly delineates where partial communities can be feasible, especially when certain components of ***s*** are disproportionately large or small (Section 8 of S1 Text). In this sense, our framework provides a static geometric characterization of the mapping from possible capture rates (***s***) to candidate communities. A natural next step is therefore to add explicit dynamics to this geometric picture: when these ***s*** vary, community transitions would correspond to trajectories crossing boundaries between feasibility partitions—moving toward larger communities (assembly) or retreating to smaller ones (disassembly). Previous work has shown how such transitions can be modeled once the relevant partition structure is specified [44, 45]; our contribution here is to define the energy-related structure on which such dynamics would unfold.

Another unresolved question is how total energy supply (*Q*), or changes in it, are translated into changes in capture rates ***s*** and thereby drive such dynamics. Currently, the decline in feasibility with larger *Q* should not be readily interpreted as a direct mechanistic prediction of collapse. Rather, it quantifies a coarse-grained constraint by measuring relevant ***s*** at fixed *Q*. Addressing this question more mechanistically would require an explicit bridge from *Q* to realized ***s***, likely through an explicit consumer-resource network in which resource supply, uptake, and cross-feeding determine effective capture rates. This perspective has motivated work on how external energy fluxes and metabolic constraints shape the diversity and structure in microbial systems [21, 22], which is also the setting where abundant free energy may be most relevant empirically [46]. In that sense, our framework can be viewed as a complementary “energetic envelope”: a coarse-grained constraint within which more detailed dynamics unfold.

In summary, our work offers a formal framework for integrating energy flux and constraints into the feasibility of complex living systems. By quantifying the energetic boundaries of ecosystem development, we provide a foundation for understanding the physical limits within which life can emerge and persist. While this framework can be extended towards more mechanistic ecological settings, it should be understood as a baseline description, built on a simplified energetic representation. Nevertheless, our findings suggest that the persistence of complex ecological communities may depend not on maximizing energy supply, but on remaining within a bounded energetic regime. By providing a formal baseline to characterize these energetic regimes, our work sets the stage for future empirical investigations to bridge specific ecological observations with these broader organizational principles.

## Supporting information

**S1 Text. Supplementary Methods and Analyses**. Detailed mathematical derivations, validation of the volume estimation pipeline, functional data analysis procedures, and sensitivity analyses across broader parameter spaces.

## Acknowledgements

We thank Daniel Rothman for insightful discussions. We acknowledge the MIT Office of Research Computing and Data for providing high performance computing resources that have contributed to the research results reported within this paper.

