## Supplemental Information for "Energetic constraints shape the diversity of feasible ecological networks"

### Table of Contents

**S1 Geometry of feasibility domains** **3**

**S2 Computing feasibility domains** **5**

**S3 Functional analysis on probabilities** **12**

**S4 Sensitivity analysis on network parameters** **15**

**S5 The critical and optimal energy supply** **21**

**S6 Partially supplied ecological networks** **23**

**S7 Density-dependent capture rates** **24**

**S8 Extension to feasibility partitions** **27**

### S1 Geometry of feasibility domains

For an $S$-species ecological network with energy demands $\boldsymbol{d}$, energy exchange $\boldsymbol{\sigma}$, total energy supply $Q$ and minimal biomass $\boldsymbol{N}^0$, its initialization domain is defined as

$$D_I(\boldsymbol{\sigma}, \boldsymbol{d}|Q, \boldsymbol{N}^0) = \{\boldsymbol{s} \in \mathbb{R}^S | \boldsymbol{s} \geq \boldsymbol{0},\ \boldsymbol{s}^T \boldsymbol{N}^0 \leq Q,\ \boldsymbol{\sigma}^{-1}(\boldsymbol{s} - \boldsymbol{d}) > \boldsymbol{0}\}, \quad (1)$$

and its maturation domain is defined as

$$D_M(\boldsymbol{\sigma}, \boldsymbol{d}|Q, \boldsymbol{N}^0) = \{\boldsymbol{s} \in \mathbb{R}^S | \boldsymbol{s} \geq \boldsymbol{0},\ \boldsymbol{s}^T \boldsymbol{N}^0 \leq Q,\ \boldsymbol{\sigma}^{-1}(\boldsymbol{s} - \boldsymbol{d}) > \boldsymbol{0}, \boldsymbol{s}^T \boldsymbol{\sigma}^{-1}(\boldsymbol{s} - \boldsymbol{d}) \leq Q\}. \quad (2)$$

#### Geometry of the initialization domain

Note that the initialization domain is a convex polytope or an empty set. Therefore, when it is feasible, standard computational geometry algorithms are available for initialization domain and remain effective in high dimensions. We will discuss the computation methods in Section S2.

*Statement* 1. The initialization domain $D_I(\boldsymbol{\sigma}, \boldsymbol{d}|Q, \boldsymbol{N}^0)$ is a *convex polytope.*

*Proof.* We can unpack the definition of $D_I(\boldsymbol{\sigma}, \boldsymbol{d}|Q, \boldsymbol{N}^0)$ as the union of $(2S + 1)$ linear constraints:

$$\boldsymbol{s} \in D_I(\boldsymbol{\sigma}, \boldsymbol{d}|Q, \boldsymbol{N}^0) \Leftrightarrow \begin{cases} s_i \geq 0, & \forall\ 1 \leq i \leq S \\ \sum_j \Lambda_{ij}(s_j - d_j) > 0, & \forall\ 1 \leq i \leq S \\ \sum_j s_j N_j^0 \leq Q. & \end{cases} \quad (3)$$

Moreover, the first and third condition(s) is equivalent to $\boldsymbol{s} \in D_C(Q, \boldsymbol{N}^0)$ (the capture domain), which is a finite simplex region. As its subset, $D_I(\boldsymbol{\sigma}, \boldsymbol{d}|Q, \boldsymbol{N}^0)$ is also a bounded subset of $\mathbb{R}^S$, if it is feasible. Eventually, $D_I(\boldsymbol{\sigma}, \boldsymbol{d}|Q, \boldsymbol{N}^0)$ is a convex polytope, if not an empty set. ∎

#### Convexity condition for the maturation domain

We assume throughout this project that $\boldsymbol{\sigma}$ is *Volterra dissipative* [1], meaning that $\frac{1}{2}(\boldsymbol{\sigma} + \boldsymbol{\sigma}^T)$ is positive definite. This restriction is an intentional modeling choice: it ensures global stability of the underlying Lotka–Volterra dynamics [2], allowing us to focus on feasibility without simultaneously treating the stability problem. A useful mathematical consequence is that it also implies convexity of the maturation domain $D^M(\boldsymbol{\sigma}, \boldsymbol{d}|Q, \boldsymbol{N}^0)$, which greatly simplifies the geometric analysis.

*Statement* 2. The maturation domain $D^M(\boldsymbol{\sigma}, \boldsymbol{d}|Q, \boldsymbol{N}^0)$ is convex if $\boldsymbol{\sigma}$ is Volterra dissipative.

*Proof.* Let us denote $H(\boldsymbol{s}; \boldsymbol{\sigma}, \boldsymbol{d}) = \boldsymbol{s}^T \boldsymbol{\sigma}^{-1}(\boldsymbol{s} - \boldsymbol{d})$ the total energy capture at steady state, as a function of $\boldsymbol{s}$, it has the quadratic component $h(\boldsymbol{s}; \boldsymbol{\sigma}, \boldsymbol{d}) = \boldsymbol{s}^T \boldsymbol{\sigma}^{-1} \boldsymbol{s}$. Hence, $h(\boldsymbol{s}; \boldsymbol{\sigma}, \boldsymbol{d}) \equiv \boldsymbol{s}^T \boldsymbol{P} \boldsymbol{s}$, where $\boldsymbol{P} = \frac{1}{2}(\boldsymbol{\sigma}^{-1} + (\boldsymbol{\sigma}^{-1})^T)$ is the symmetrization of $\boldsymbol{\sigma}^{-1}$.

Since $\boldsymbol{\sigma}$ is invertible, any given $\boldsymbol{s} \in \mathbb{R}^S$ is uniquely associated with $\boldsymbol{x} = \boldsymbol{\sigma}^{-1}\boldsymbol{s}$. Since $\boldsymbol{\sigma}$ is Volterra dissipative, for any $\boldsymbol{s}$, $h(\boldsymbol{s}; \boldsymbol{\sigma}, \boldsymbol{d}) = (\boldsymbol{\sigma x})^T \boldsymbol{P} (\boldsymbol{\sigma x}) = \boldsymbol{x}^T (\boldsymbol{\sigma}^T \boldsymbol{P} \boldsymbol{\sigma}) \boldsymbol{x} = \boldsymbol{x}^T (\frac{1}{2}\boldsymbol{\sigma}^T + \frac{1}{2}\boldsymbol{\sigma}) x > 0$. Therefore, $h(\boldsymbol{s}; \boldsymbol{\sigma}, \boldsymbol{d}) = \boldsymbol{s}^T \boldsymbol{P} \boldsymbol{s}$ is a positive definite quadratic form.

For $H(\boldsymbol{s}) = h(\boldsymbol{s}) - \boldsymbol{s}^T \boldsymbol{\sigma}^{-1} \boldsymbol{d}$, the linear term $-\boldsymbol{s}^T \boldsymbol{\sigma}^{-1} \boldsymbol{d}$ does not change the convexity of $H$ function. Therefore, The sub-level set of $H$, i.e., $\{\boldsymbol{s}|H(\boldsymbol{s}) \leq Q\}$ is a convex domain [3]. Finally, as the intersection of two convex domains, the maturation domain $D_M(\boldsymbol{\sigma}, \boldsymbol{d}; Q, \boldsymbol{N}^0) = D_I(\boldsymbol{\sigma}, \boldsymbol{d}; Q, \boldsymbol{N}^0) \cap \{\boldsymbol{s}|H(\boldsymbol{s}) \leq Q\}$ is also convex. ■

#### Geometry of the maturation domain

The maturation domain is the intersection of the initialization domain and the sub-level set of function $H(\boldsymbol{s})$. We have shown that $H(\boldsymbol{s})$ is convex under certain conditions, here we further illustrate that $\{\boldsymbol{s}|H(\boldsymbol{s}) \leq Q\}$ becomes an ellipsoid.

*Statement* 3. $\{\boldsymbol{s}|H(\boldsymbol{s}) \leq Q\}$ defines an *ellipsoid* in $\mathbb{R}^S$ if $\boldsymbol{\sigma}$ is Volterra dissipative.

*Proof.* Notice that $\boldsymbol{P}$ is a positive definite symmetrical matrix, which can be *Cholesky decomposed* as $\boldsymbol{P} = \boldsymbol{L}^T \boldsymbol{L}$ [4]. Let us denote $\boldsymbol{c} = -\boldsymbol{\sigma}^{-1}\boldsymbol{d}$, $\boldsymbol{y} = \boldsymbol{L s}$, we complete the square in $H$:

$$\begin{aligned} H(\boldsymbol{s}) &= \boldsymbol{s}^T \boldsymbol{\sigma}^{-1}(\boldsymbol{s} - \boldsymbol{d}) = \boldsymbol{s}^T \boldsymbol{L}^T \boldsymbol{L} \boldsymbol{s} + \boldsymbol{c}^T \boldsymbol{s} \\ &= \boldsymbol{y}^T \boldsymbol{y} + \boldsymbol{c}^T \boldsymbol{L}^{-1} \boldsymbol{y} \\ &= \underbrace{\left(\boldsymbol{y} + \frac{1}{2}(\boldsymbol{L}^T)^{-1}\boldsymbol{c}\right)^T \left(\boldsymbol{y} + \frac{1}{2}(L^T)^{-1}\boldsymbol{c}\right)}_{\triangleq \tilde{h}(\boldsymbol{y})} - \frac{1}{4}\boldsymbol{c}^T \boldsymbol{P}^{-1} \boldsymbol{c}. \end{aligned} \tag{4}$$

Therefore, $H(\boldsymbol{s}) \leq Q \Leftrightarrow \tilde{h}(\boldsymbol{y}) \leq Q + \frac{1}{4}\boldsymbol{c}^T \boldsymbol{P}^{-1}\boldsymbol{c} = t$, which defines a *ball* in the transformed space of $\boldsymbol{y}$. This ball has its center at $\boldsymbol{y}_c = -\frac{1}{2}(\boldsymbol{L}^T)^{-1}\boldsymbol{c}$ and square of radius $t = Q + \frac{1}{4}\boldsymbol{c}^T \boldsymbol{P}^{-1}\boldsymbol{c}$. Since $\boldsymbol{y} = \boldsymbol{L s}$, $\{H(\boldsymbol{s}) \leq Q\}$ is $\{\tilde{h}(\boldsymbol{y}) \leq t\}$ linear transformed via $\boldsymbol{L}^{-1}$, which is an ellipsoid.

■

As we shall see in the next section, the above analysis has motivated us to study $D^M(\boldsymbol{\sigma}, \boldsymbol{d}|Q, \boldsymbol{N}^0)$ at the transformed space of $\boldsymbol{y}$, where the quadratic constraint becomes a standard ball constraint.

### S2 Computing feasibility domains

In this section, we discuss the methods to perform computational geometry analysis on both the initialization and maturation domains. We present the pipeline to 1) standardize these tasks into a unified mathematical form, 2) perform sampling and volume estimation on the standard form and 3) translate these numerical results back into the original problem. Along with this document, we provide complete *Julia* code implementation of these methods.

#### Exact computation of the initialization domain

First of all, $\text{vol}(D_I(\boldsymbol{\sigma}, \boldsymbol{d}|Q, \boldsymbol{N}^0))$ can be computed *exactly* and efficiently. As a well-established problem in computational geometry, the standard method decomposes the polytope into simplices and sum up their volumes that are given analytically [5]. This method is realized inside the Polyhedra package in *Julia* [6], whose lower-end realization relied on the widely used *Qhull* algorithm [7]. We utilize these available repositories to calculate the volume of $D_I(\boldsymbol{\sigma}, \boldsymbol{d}|Q, \boldsymbol{N}^0))$. In what follows, we discuss the *approximation* methods based on uniform sampling, which applies to all the domains defined in this project.

#### The *InterPolyBalls* class

As we see in the previous section, the geometrical complexity of $D^{C,I,M}(\boldsymbol{\sigma}, \boldsymbol{d}|Q, \boldsymbol{N}^0)$ remains unchanged for any set of parameters $\{\boldsymbol{\sigma}, \boldsymbol{d}, Q, \boldsymbol{N}^0\}$. Moreover, we could standardize these domains into a unified mathematical form, which we call the *InterPolyBalls* class.

An *InterPolyBalls* object $D(A, \boldsymbol{b}, \{(\boldsymbol{c}_i, t_i)\})$ is the enclosed intersecting convex body of:

1) A set of linear constraints (a polytope): $\boldsymbol{Ax} \leq \boldsymbol{b}$;
2) A set of ball constraints: $(\boldsymbol{x} - \boldsymbol{c}_i)^T(\boldsymbol{x} - \boldsymbol{c}_i) \leq t_i$.

To simplify the notation, we again denote $\boldsymbol{c} = -\boldsymbol{\sigma}^{-1}\boldsymbol{d}$, $\boldsymbol{P} = \frac{1}{2}(\boldsymbol{\sigma}^{-1} + (\boldsymbol{\sigma}^{-1})^T)$ and $\boldsymbol{L}^T\boldsymbol{L} = \boldsymbol{P}$.

Next, we derive how to standardize feasibility domains as the unified *InterPolyBalls* class. Then, we develop computational methods on the *InterPolyBalls* class.

#### Feasibility domains as *InterPolyBalls*

As we can see directly, the initialization domain is a special case of *InterPolyBalls* without ball constraints.

*Statement* 4. $D_I(\boldsymbol{\sigma}, \boldsymbol{d}|Q, \boldsymbol{N}^0)$ is an *InterPolyBalls*.

*Proof.* We note that $D_I(\boldsymbol{\sigma}, \boldsymbol{d}|Q, \boldsymbol{N}^0)$ is equivalent to $D(\boldsymbol{A}_0, \boldsymbol{b}_0, \{\})$, with the following parameters:

$$\boldsymbol{A_0} = \begin{pmatrix} -I \\ -\boldsymbol{\sigma}^{-1} \\ \boldsymbol{N}^{0T} \end{pmatrix}, \ \boldsymbol{b}_0 = \begin{pmatrix} \boldsymbol{0} \\ \boldsymbol{c} \\ Q \end{pmatrix}, \tag{5}$$

since $\boldsymbol{A}_0\boldsymbol{s} \leq \boldsymbol{b}_0$ completely represent the polytope defining $D_I(\boldsymbol{\sigma}, \boldsymbol{d}|Q, \boldsymbol{N}^0)$, and $\{\}$ indicates there is no quadratic constraint. ■

Second, as we see in the previous section, the additional constraint for the maturation domain is an ellipsoid body, which could be linear transformed as a ball constraint. This allows us to standardize the maturation domain.

*Statement* 5. $D_M(\boldsymbol{\sigma}, \boldsymbol{d}|Q, \boldsymbol{N}^0)$ can be represented as an *InterPolyBalls* up to the linear transformation given by $\boldsymbol{L}$.

*Proof.* Given the $\boldsymbol{\sigma}$ matrix in $D_M(\boldsymbol{\sigma}, \boldsymbol{d}|Q, \boldsymbol{N}^0)$, which is assumed to be Volterra dissipative, we construct the corresponding $\boldsymbol{L}$. According to Eq. (4), $H(\boldsymbol{s}) \leq Q$ is equivalent to $\tilde{h}(\boldsymbol{L}^{-1}\boldsymbol{y}) \leq t$, and $\tilde{h}$ defines a standard ball. This motivate us to linear transform $D_M$ into $\tilde{D_M} = \boldsymbol{L}\,(D_M)$.

For any $\boldsymbol{y} \in \tilde{D_M}(\boldsymbol{\sigma}, \boldsymbol{d}|Q, \boldsymbol{N}^0)$, its *preimage* $\boldsymbol{s} = \boldsymbol{L}^{-1}\boldsymbol{y}$ satisfies

$$\begin{cases} \boldsymbol{L}^{-1}\boldsymbol{y} & \geq \boldsymbol{0} \\ \boldsymbol{\sigma}^{-1}\boldsymbol{L}^{-1}\boldsymbol{y} & > \boldsymbol{\sigma}^{-1}\boldsymbol{d} \\ {\boldsymbol{N}^0}^T\boldsymbol{L}^{-1}\boldsymbol{y} & \leq Q \\ \boldsymbol{y}^T\boldsymbol{y} + \boldsymbol{c}^T\boldsymbol{L}^{-1}\boldsymbol{y} & \leq Q \end{cases}, \tag{6}$$

therefore, $\tilde{D_M}(\boldsymbol{\sigma}, \boldsymbol{d}|Q, \boldsymbol{N}^0)$ is an *InterPolyBalls* object $D(\boldsymbol{A}_1, \boldsymbol{b}_1, \{(\boldsymbol{y}_c, t)\})$, whose parameters read

$$\boldsymbol{A}_1 = \begin{pmatrix} -\boldsymbol{L}^{-1} \\ -\boldsymbol{\sigma}^{-1}\boldsymbol{L}^{-1} \\ {\boldsymbol{N}^0}^T\boldsymbol{L}^{-1} \end{pmatrix}, \ \boldsymbol{b}_1 = \begin{pmatrix} \boldsymbol{0} \\ \boldsymbol{c} \\ Q \end{pmatrix}, \ \boldsymbol{y}_c = -\frac{1}{2}(\boldsymbol{L}^{-1})^T\boldsymbol{c}, \ t = Q + \frac{1}{4}\boldsymbol{c}^T\boldsymbol{P}^{-1}\boldsymbol{c}. \tag{7}$$

The above standardization invokes linear transformation, which will be compatible with our computational framework of sampling and volume estimation. ■

Next, we discuss the pipeline to perform Markov chain Monte Carlo sampling and associated stochastic volume estimation on any*InterPolyBalls* domain $D(A, \boldsymbol{b}, \{(\boldsymbol{c}_i, t_i)\})$.

#### Pre-processing on *InterPolyBalls*

As a standard starting step, pre-processing could help to find a proper starting point of sampling, provide reference for volume estimation, and enhance the performance of the computational methods. Here, we adopt pre-processing by first solving the maximal inscribed ball (chevbyshev ball) problem for $D(\boldsymbol{A}, \boldsymbol{b}, \{(\boldsymbol{c}_i, t_i)\}$. The outcome, $\mathbb{B}(\boldsymbol{x}^*, r^*)$, will be important for further sampling and volume estimation. For the *InterPolyBalls* class, this step requires optimization over second-order cone constraints and linear constraints (see Alg. 1). Our implementation of pre-processing can be found in the `chevball` function of the code repository.

**Algorithm 1:** Compute Chebyshev Ball of *InterPolyBalls* Domain

**Input:** *InterPolyBalls* Domain $D(\boldsymbol{A}, \boldsymbol{b}, \{(\boldsymbol{c}_i, t_i)\}$

**Output:** Largest inscribed ball $\mathbb{B}(\boldsymbol{x}^*, r^*) \subseteq D$

Define $\|A\|_{\text{row},2}$ the vector of 2-norm of $A$'s rows

**Solve the following convex optimization problem** $p$**:**

$$\begin{aligned} \text{maximize} \quad & r \\ \text{subject to} \quad & A\boldsymbol{x} + r \cdot \|A\|_{\text{row},2} \leq \boldsymbol{b} \\ & \|x - \boldsymbol{c}_i\|_2 \leq \sqrt{t_i} - r \\ & r \geq 0 \end{aligned}$$

**if** *p has a feasible optimal solution* **then**
  **return** $\mathbb{B}(\boldsymbol{x}^*, r^*)$
**else**
  **return** invalid flag

#### Sampling on *InterPolyBalls*

A standard step in computing geometrical objects like *InterPolyBalls* is to uniformly sample within the domain. Namely, generating a sequence of samples $\{\boldsymbol{y}_i\}$ from the distribution $\mathbf{y} \sim \texttt{Unif}(D(A, \boldsymbol{b}, \{(\boldsymbol{c}_i, t_i)\})$. As the general approach, *Markov Chain Monte Carlo* (MCMC) sampling is widely used for this kind of tasks. The basic idea is to first find a proper starting point $\boldsymbol{y}_0 \in D(A, \boldsymbol{b}, \{(\boldsymbol{c}_i, t_i)\})$, and then perform a designed random walk (Markov Chain), where the trajectories $\{\boldsymbol{y}_t\}$ converge to a sequence of uniform samples.

In the context of MCMC on a $S$-dimensional convex body $D$, *hit-and-run sampling* proves to be the effective algorithm [8–11]. It performs random walks by repeating the following procedures:

**Algorithm 2:** Hit-and-Run MCMC on Convex Body $D \subset \mathbb{R}^S$

**Input:** Initial point $\boldsymbol{y}_0 \in D$, number of steps $T$

**Output:** Sequence $\{\boldsymbol{y}_t\}_{t=0}^T \subset D$

**for** $t = 0$ **to** $T - 1$ **do**
  Sample random direction $\boldsymbol{\Delta}_t \in \mathbb{R}^S$;
  Compute bounds $\lambda^-, \lambda^+$ such that $\boldsymbol{y}_t + \lambda\boldsymbol{\Delta}_t \in D$ for $\lambda \in [\lambda^-, \lambda^+]$;
  Sample $\lambda_t \sim \texttt{Unif}[\lambda^-, \lambda^+]$;
  Set $\boldsymbol{y}_{t+1} \leftarrow \boldsymbol{y}_t + \lambda_t\boldsymbol{\Delta}_t$;

In relate to the *InterPolyBalls* class, the bounds $(\lambda^-, \lambda^+)$ are given by examination each constraint (boundary) in $D(A, \boldsymbol{b}, \{(\boldsymbol{c}_i, t_i)\})$. Specifically, each linear constraint (half-space) intersecting with $\boldsymbol{y}_t + \lambda\boldsymbol{\Delta}_t$ line at one point, contributing to one bound in $\lambda$ (upper or lower); each quadratic constraint (ball) intersecting with $\boldsymbol{y}_t + \lambda\boldsymbol{\Delta}_t$ line at two points, contributing to two bounds in $\lambda$ (upper and lower). Practically, computing these bounds is equivalent to solving the critical $\lambda$ values at which the $\boldsymbol{y}_t + \lambda\boldsymbol{\Delta}_t$ line intersects with $D$ through the following equations:

$$\begin{cases} (A\boldsymbol{y}_t)_j + \lambda(A\boldsymbol{\Delta}_t)_j = b_j \\ (\boldsymbol{y}_t + \lambda\boldsymbol{\Delta} - \boldsymbol{c}_i)^T(\boldsymbol{y}_t + \lambda\boldsymbol{\Delta} - \boldsymbol{c}_i) = t_i \end{cases} \tag{8}$$

The final bounds are given by the filtered extrema of the list of critical values $\boldsymbol{\lambda}$, i.e. $\lambda^+ := \min(\{\lambda_i \in \boldsymbol{\lambda} \mid \lambda_i > 0\} \cup \{0\})$ and $\lambda^- := \max(\{\lambda_i \in \boldsymbol{\lambda} \mid \lambda_i < 0\} \cup \{0\})$. The code to implement a single step in Alg. 2 and the bounds computing can be found in the `hr_step` function.

To generate quality samples, we could leverage warm-up techniques and multi-threads sampling. For warm-up, we uniformly select a point inside the Chebyshev ball $\mathbb{B}(\boldsymbol{x}^*, r^*)$) of the domain as the initial point to perform Alg. 2, i.e. $\boldsymbol{y}_0 \sim \texttt{Unif}(\mathbb{B}(\boldsymbol{x}^*, r^*))$. The motivation for such choice is that, $\boldsymbol{y}_0$ is guaranteed to be inside $D$, which avoids algorithm failure; $\boldsymbol{y}_0$ is not too close to any boundary of $D$, which enhance the chance of fast convergence towards uniform sampling [12]. For multi-threads sampling, we restart the random walk with a new $\boldsymbol{y}_0$ and repeat Alg. 2 for several times (threads), which allows for convergence diagnosis and provides robustness from a "bad" starting point. In using these methods for our simulated possibility domains, we generate $2 \times 10^4 - 2 \times 10^5$ samples per thread, and we keep only the second half of each thread (burn-in step); we repeat the sampling for 10 threads. In total, we get $10^5 - 10^6$ samples per domain, which is tested to provide robust and uniform representations of the possibility domain. This level of sampling ensures statistical reliability in further estimating volume ratios. The code to implement these designs for an entire sampling task can be found in the `hr_sample` function.

#### Volume estimation of *InterPolyBalls*

We implement the *Multiphase Monte Carlo* method to estimate the volume of *InterPolyBalls*. We note that similar approaches have recently shown success in [13, 14]. The overall idea of this method is to construct a series of convex bodies $\{B_i\}$ between the focal domain ($D$) and an analytically computable domain ($\mathbb{B}(\boldsymbol{x}^*, r^*)$), such that $\mathbb{B}(\boldsymbol{x}^*, r^*) = B_0 \subset B_1 \subset \ldots B_n = D$. Then, estimating the volume of $D$ through

$$\underbrace{\mathrm{vol}(B_n)}_{=\mathrm{vol}(D)} = \underbrace{\mathrm{vol}(B_0)}_{\text{ball}} \prod_{i=0}^{n-1} \frac{\mathrm{vol}(B_{i+1})}{\mathrm{vol}(B_i)}, \tag{9}$$

where $\mathrm{vol}(B_0) = \dfrac{\pi^{\frac{S}{2}} r^{*S}}{\Gamma(\frac{S}{2}+1)}$ is analytically computable, and ratio $\dfrac{\mathrm{vol}(B_{i+1})}{\mathrm{vol}(B_i)}$ can be estimated by counting the ratio of uniform samples from $B_{i+1}$ that falls into $B_i$. Namely, for $\mathsf{y} \sim \texttt{Unif}(B_{i+1})$, $\dfrac{\mathrm{vol}(B_i)}{\mathrm{vol}(B_{i+1})} = \mathbb{E}(\mathbb{1}_{\mathsf{y} \in B_i})$.

We construct such $B_i$ in the following way. We first run a small sampling over domain $D$ (in total $5 \times 10^3$ samples), then find the largest distance ($\rho$) between each sample and center of the Chebyshev ball $\boldsymbol{x}^*$. $\rho$ measures approximately the largest distance between $\boldsymbol{x}^*$ and any point in $D$. With high probability, $D = B_n \subset \mathbb{B}(\boldsymbol{x}^*, \rho)$, and thus $D = D \cap \mathbb{B}(\boldsymbol{x}^*, \rho') = B_{\rho'}$, for sufficiently large $\rho' > \rho$.

Therefore, we can set $B_i = D \cap \mathbb{B}(\boldsymbol{x}^*, \rho_i)$, where $\rho_i = r(\frac{\rho}{r})^{\frac{i}{N}}$. Collectively, $\rho_i, 0 \leq i \leq n$ creates a set of concentric balls $\{\mathbb{B}(\boldsymbol{x}^*, \rho_i)\}$, and $B_i$ is created by each of these balls intersecting with $D$. Notice that the first ball $i = 0, \rho_i = r$ is just the Chebyshev ball of $D$; the last few balls $N \leq i \leq n$ enclose $D$, thus $B_i = D$. By definition, $B_i$ is still an *InterPolyBalls* object, which means their volume ratios can be estimated by using the previously discussed sampling methods. In our current implementation, we set $N = 10, n = 12$. For more details, refer to the `volume_domain` function.

#### Volume estimation for feasibility domains

Particularly, we take the following approach to simply computation for computing $\mathbb{P}^{I,M}(Q)$ over a range of $Q \in [Q_{\min}, Q_{\max}]$. Instead of independently measuring each $D(Q)$ that is related to $D_{I,\tilde{M}}(\boldsymbol{\sigma}, \boldsymbol{d}|Q, \boldsymbol{N}^0)$, we could instead leverage multiphase Monte Carlo over the set of domains $\{D(Q), Q \in [Q_{\min}, Q_{\max}]\}$. Specifically, notice that $D_{I,\tilde{M}}(\boldsymbol{\sigma}, \boldsymbol{d}|Q_i, \boldsymbol{N}^0) \subset D_{I,M}(\boldsymbol{\sigma}, \boldsymbol{d}|Q_{i+1}, \boldsymbol{N}^0)$, for $Q_{i+1} > Q_i$. Linear transformation preserves the subset relationship, thus $D(Q_i) \subset D(Q_{i+1})$. Then, we could directly estimate $\dfrac{\mathrm{vol}(D(Q_i))}{\mathrm{vol}(D(Q_{i+1}))}$ by sampling uniformly from $D(Q_{i+1})$ and compute the ratio of samples that fall into $D(Q_i)$. Finally, we execute the original volume estimation once for $\mathrm{vol}(D(Q_{\max}))$, the largest domain, and then we estimate $\mathrm{vol}(D(Q))$ as:

$$\mathrm{vol}(D(Q)) = \mathrm{vol}(D(Q_{\max})) \prod_{k=n-1}^{i} \frac{\mathrm{vol}(D(Q_k)}{\mathrm{vol}(D(Q_{k+1})}, \tag{10}$$

where $Q_n = Q_{\max}$ and $Q_i = Q$. The error for estimating $\mathrm{vol}(D(Q))$ is acceptable when there are enough samples for each ratio estimation, and when the interval $\Delta Q = Q_{i+1} - Q_i$ is small enough. In this way, we could save the intermediate $N$ samplings for other $Q$ values and reduce computation time significantly. For more details, refer to the `volume_range_EFD` function.

#### Translate back into original domains

Finally, we discuss how to sample and measure the volumes of feasibility domains.

The initialization domain $D_I(\boldsymbol{\sigma}, \boldsymbol{d}|Q, \boldsymbol{N}^0)$ is itself an *InterPolyBalls* object with parameters given in (5). Therefore, we can directly apply the sampling methods in section S2.5 to study the properties of $\boldsymbol{s} \in D_I(\boldsymbol{\sigma}, \boldsymbol{d}|Q, \boldsymbol{N}^0)$. Conveniently, we use the exact volume formula in S1.1 to measure $\mathrm{vol}(D_I(\boldsymbol{\sigma}, \boldsymbol{d}|Q, \boldsymbol{N}^0))$.

The maturation domain $D_M(\boldsymbol{\sigma}, \boldsymbol{d}|Q, \boldsymbol{N}^0)$ is an *InterPolyBalls* subject to a linear transformation. Once we apply sampling and volume estimation on the transformed domain $\tilde{D_M}$ (defined and parameterized in (7)), the following relationships instantly translate the volume estimation and sampling results back to $D_M$.

*Statement* 6. The volumes follows $\mathrm{vol}(D_M) = \mathrm{vol}(\tilde{D_M}) / \det(\boldsymbol{L})$

*Proof.*

$$\begin{aligned} \mathrm{vol}(D_M) &= \int_{D_M} \mathrm{d}\boldsymbol{s} \\ &= \int_{\tilde{D_M}} \mathrm{d}\boldsymbol{y} \left|\frac{\mathrm{d}\boldsymbol{s}}{\mathrm{d}\boldsymbol{y}}\right| \qquad (\boldsymbol{y} = \boldsymbol{L}\boldsymbol{s}; \tilde{D_M} = \boldsymbol{L}(D_M)) \\ &= \det(\boldsymbol{L}^{-1}) \int_{\tilde{D_M}} \mathrm{d}\boldsymbol{y} \qquad (\text{Jacobian } \frac{\mathrm{d}\boldsymbol{s}}{\mathrm{d}\boldsymbol{y}} = \boldsymbol{L}^{-1}) \\ &= \mathrm{vol}(\tilde{D_M}) / \det(\boldsymbol{L}). \end{aligned} \tag{11}$$

■

*Statement* 7. The uniform samples hold up to the same linear transformation $\boldsymbol{L}$

*Proof.* For $\mathsf{y} \sim \texttt{Unif}(\tilde{D_M})$, i.e. $p_{\mathsf{y}}(\boldsymbol{y}) = \begin{cases} \frac{1}{\mathrm{vol}(\tilde{D_M})} & \boldsymbol{y} \in \tilde{D_M} \\ 0 & \text{otherwise} \end{cases}$, the random variable defined from function $\mathsf{s} = \boldsymbol{L}^{-1}\mathsf{y}$ is associated by

$$\int p_{\mathsf{s}}(\boldsymbol{s})\mathrm{d}\boldsymbol{s} = \int p_{\mathsf{y}}(\boldsymbol{y}(\boldsymbol{s}))\mathrm{d}\boldsymbol{y}. \tag{12}$$

Therefore, the probability distribution function for $\mathsf{s}$ reads

$$p_{\mathsf{s}}(\boldsymbol{s}) = p_{\mathsf{y}}(\boldsymbol{y}(\boldsymbol{s}))\left|\frac{\mathrm{d}\boldsymbol{y}}{\mathrm{d}\boldsymbol{s}}\right| = \begin{cases} \dfrac{1}{\mathrm{vol}(\tilde{D_M})}\det(\boldsymbol{L}) & \boldsymbol{y}(\boldsymbol{s}) \in \tilde{D_M} \\ 0 & \text{otherwise} \end{cases} = \begin{cases} \dfrac{1}{\mathrm{vol}(D_M)} & \boldsymbol{s} \in D_R \\ 0 & \text{otherwise} \end{cases}. \tag{13}$$

Hence $\mathsf{s}$ is the uniform distribution on $D_M$. ∎

Therefore, sampling $\mathsf{y}$ from $\tilde{D_M}$ and transform them via $\boldsymbol{s} = \boldsymbol{L}^{-1}\boldsymbol{y}$ gives uniform sampling on $D_M$; and estimating the volume of $\tilde{D_M}$ directly gives the estimation of volume of $D_M$.

#### Validate the computation

In the end, we validate our computational pipeline with test cases. We consider measuring volume of domain $D(S, Q) = \{\boldsymbol{s} \in \mathbb{R}^S | \boldsymbol{s} \geq \boldsymbol{0},\ \boldsymbol{s}^\top \boldsymbol{s} \leq r^2\}$,which is the positive orthant of a ball in dimension $S$ with radius $r$. We compose the test cases using $S \in [2, 4, 8, 12]$ and $r \in [10^0, 10^2]$ that is logarithmically spaced into 25 values. Then we calculate their volumes independently using 10 threads and $10^4$ samples in each thread (in total $5 \times 10^4$ effective samples). Each domain takes around 1 second to calculate on a personal computer with M2pro processor with *Julia* v1.11.3, which is highly efficient. Finally, we compare the estimations with the analytical solutions in terms of relative error. The results are shown in Fig. S1, suggesting outstanding accuracy of the computation pipeline.

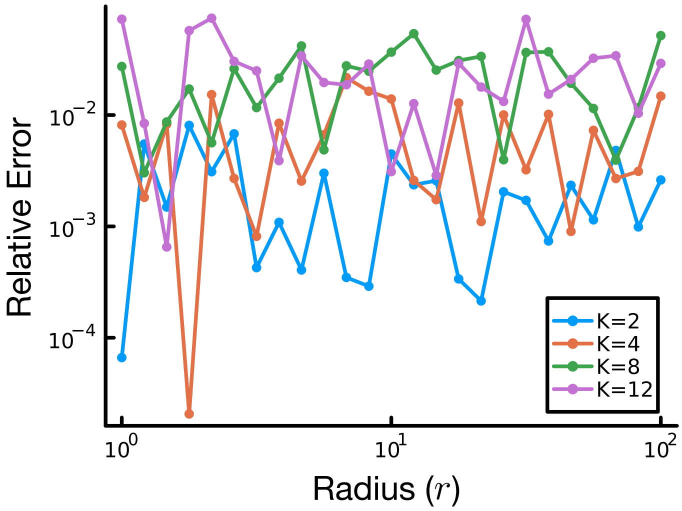


Figure S1: **Relative errors across test cases** Each colored line shows the relative error under a given dimension with increasing radius. Domains with $S = \{2, 4, 8, 12\}$ have average relative error of $\{2.2 \times 10^{-3}, 7.1 \times 10^{-3}, 2.3 \times 10^{-2}, 2.7 \times 10^{-2}\}$. Higher dimensions yield larger relative error, which could be further compensated by adjusting number of samples and replications.

### S3 Functional analysis on probabilities

To extract general trends from large-scale network ensembles with randomized parameters $\boldsymbol{\theta} = \{\boldsymbol{\sigma}, \boldsymbol{d}, \boldsymbol{N}^0\}$, we adopt a Functional Data Analysis (FDA) approach. For each realization, the relationship between feasibility probability and energy supply Q is treated as a continuous random function $\mathbb{P}(Q; \boldsymbol{\theta})$.

#### Statistical models of probabilities

Since probabilities are strictly bounded within the interval $[0, 1]$, a simple linear or logarithmic average often leads to biased central tendencies and nonphysical confidence intervals (e.g., exceeding 1). Therefore, we introduce the following statistical model based on logit transformation.

We perform a logit transformation to map the probability space to the real line $\mathbb{R}$:

$$\lambda(Q; \boldsymbol{\theta}) = \operatorname{logit}(\mathbb{P}(Q; \boldsymbol{\theta})) = \log\left(\frac{\mathbb{P}(Q; \boldsymbol{\theta})}{1 - \mathbb{P}(Q; \boldsymbol{\theta})}\right). \tag{14}$$

To handle the numerical singularities at the boundaries $\mathbb{P} = 0, 1$, we apply a clamping epsilon ($\epsilon = 10^{-3}$) such that $P \in [\epsilon, 1 - \epsilon]$. We assume that in the transformed logit-space, the ensemble of curves follows a distribution characterized by its functional mean $\mu_\lambda(Q)$ and standard deviation $\sigma_\lambda(Q)$:

$$\begin{aligned} \mu_\lambda(Q) &= \mathbb{E}_{\boldsymbol{\theta}}[\lambda(Q; \boldsymbol{\theta})], \\ \sigma_\lambda(Q) &= \sqrt{\mathbb{V}_{\boldsymbol{\theta}}[\lambda(Q; \boldsymbol{\theta})]}. \end{aligned} \tag{15}$$

The central trend and the variability are then mapped back to the probability space using the inverse of logistic function (sigmoid function):

$$\begin{aligned} \bar{P}(Q) &= \operatorname{sigmoid}(\mu_\lambda(Q)) = \frac{1}{1 + \exp(-\mu_\lambda(Q))}, \\ P_\pm(Q) &= \operatorname{sigmoid}(\mu_\lambda(Q) \pm \sigma_\lambda(Q)). \end{aligned} \tag{16}$$

This approach yields two significant advantages for ecological interpretation. First, the estimated mean and confidence bands are strictly constrained within $(0, 1)$; second, the resulting confidence bands are inherently asymmetric in the probability space, correctly reflecting diminishing perturbation when average trend is near 0 or 1.

#### Measuring unimodal and saturation of individual curve

To further confirm that general patterns found in the previous subsection apply to each individual case (i.e., each replication of $\mathbb{P}$-$Q$ curve), we develop unimodal score ($G_U$) and saturating score ($G_S$) to quantify the unimodal and saturation pattern in $\mathbb{P}_{I,M}(Q)$. Roughly, each curve is represented as a set of data points, and these scores calculate the weighted "ratio" of points whose first and second order difference is consistent with the unimodal and saturation pattern (Figure S2 C). As a quantity within $[0, 1]$, the higher the score, the more consistent the observed data curve with the proposed pattern. Conversely, a null model of random curve produces significantly lower score.

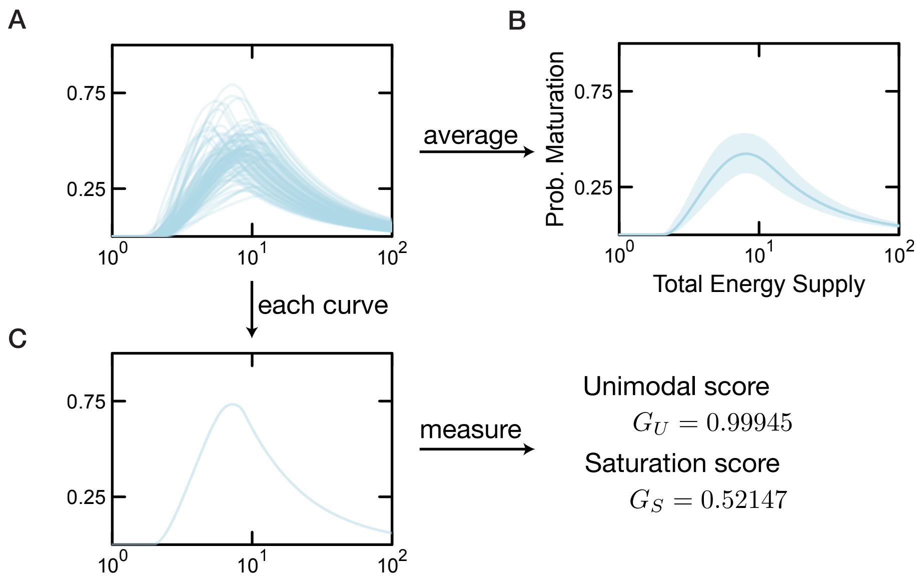


Figure S2: **Overview of the functional data analysis A** We first collect probability and total supply results from multiple cases, which show both variation and general patterns, our goal is to extract the generality and test whether it applies to each case. **B** We extract the general patterns by taking average and standard deviation at each total supply level. **C** We use the unimodal and saturation score to test if each case follow the general trend discovered.

Given a scalar function $y = f(x), x \in [x_i, x_f]$ , we define the saturation pattern as $f'(x) > 0, \forall x \in [x_i, x_f]$ and $f''(x) < 0, \forall x \in [\frac{1}{2}(x_i + x_f), x_f]$. That is, $f$ saturates in $[x_i, x_f]$ if it monotonically increase in $[x_i, x_f]$, and such increase decays for the latter half of the interval $[\frac{1}{2}(x_i + x_f), x_f]$. Notice this $1/2$ fraction is can be adjusted for different levels of sensitivity.

Similarly, the unimodal pattern is defined as $\exists ! x_c \in [x_i, x_f]$, s.t. $f'|_{x<x_c} > 0$ and $f'|_{x>x_c} < 0$, such $(x_c, f(x_c))$ naturally is the unique global maximum point for $f$.

To numerically calculate these scores, we first perform a locally estimated scatterplot smoothing (LOESS) to denoise the data points and to equalize the intervals.

The result $\{(x_i, y_i)\}_{i=1}^{N}$ is then counted at each $i = 1, \cdots, N$, where an eligible $i$ gets score proportional to $|\Delta y_i|$:

$$
\begin{aligned}
G_S &= \frac{\sum_i^{N-1} |\Delta y_i| I_i^S}{\sum_i^{N-1} |\Delta y_i|}, \quad I_i^S = I(\Delta y_i > 0) \cdot [I(x_i < \frac{1}{2}(x_i + x_f)) + I(\Delta y_{i+1} < \Delta y_i, x_i > \frac{1}{2}(x_i + x_f))] \\
G_U &= \frac{\sum_i^{N-1} |\Delta y_i| I_i^U}{\sum_i^{N-1} |\Delta y_i|}, \quad I_i^U = I(\Delta y_i > 0, x_i < \hat{x_c}) + I(\Delta y_i < 0, x_i > \hat{x_c})
\end{aligned} \tag{17}
$$

where $I$ is the indicator function. Since $I_i^S, I_i^U \in \{0, 1\}$, $G_S, G_U \in [0, 1]$.

In the main text, we showed under each $S \in \{2, 4, 6, 8\}$, the general trend generated from 100 replications, here we provide the mean of the their scores in Table S1, which are all very close to 1. In comparison, the null model that replaces these probabilities with $y_i \sim \texttt{Unif}(0, 1)$ yield much lower scores.

| Size | Unimodal score | Saturation score |
|---|---|---|
| 2 | 0.9948 | 0.9993 |
| 4 | 0.9999 | 0.9995 |
| 6 | 0.9999 | 0.9989 |
| 8 | 0.9999 | 0.9987 |
| null | 0.3261 | 0.5108 |

Table S1: Average unimodal and saturation scores for the main text results and for null model of uniformly distrbuted data points.

### S4 Sensitivity analysis on network parameters

In this section, we conduct sensitivity analysis on ecological parameters $\{\boldsymbol{\sigma}, \boldsymbol{d}, \boldsymbol{N}^0\}$ that will affect the probability of initialization and maturation. Analysis here shows a clear and interpretable change on the network's probability, yet their saturation and unimodal patterns remain highly robust.

#### Mass-specific energy demands

The mass-specific energy demands ($\boldsymbol{d}$) measures a baseline level of energy capture for each population $i$. In the main text, we set an average $d_0 = 1$. Here, we compare the it with $d_0 = 0.5$ and $d_0 = 2.0$ to mimic an increase in average demands. As seen in Fig. S3A, when the average demand level $d_0$ increases, a larger total energy supply ($Q$) is needed to compensate for this increasing demands and keep the same probability of initialization. The maximal probability of initialization at saturation, however, remains unchanged. Similarly, in Fig. S3B, the optimal total energy supply ($Q$) increases, but the overall probability of maturation decreases.

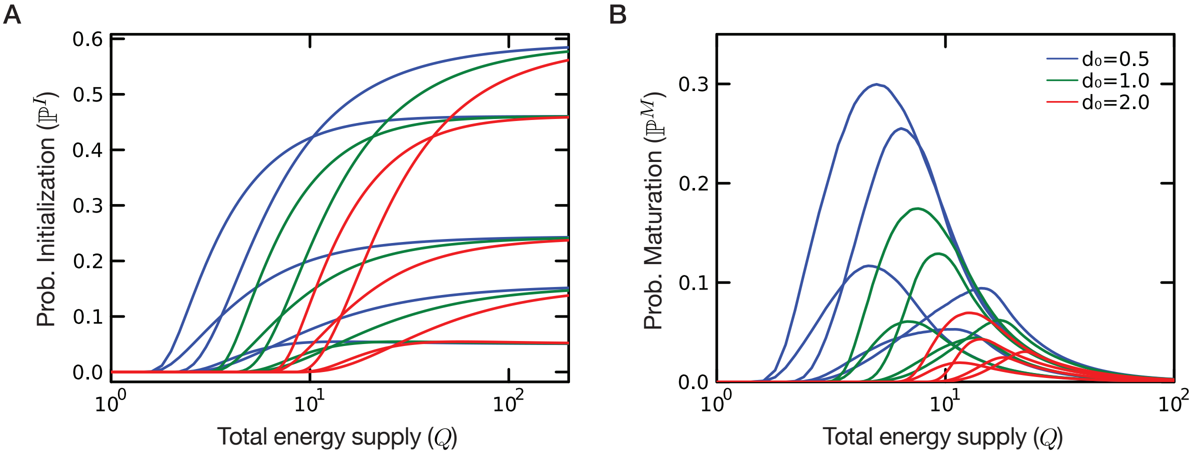


Figure S3: **Increasing average demand postpone the saturation and optimization of both probabilities, but only decrease probability of maturation.** Under $S = 4$, $N_0 = 1$, we generate five $\boldsymbol{\sigma}$ matrices and compute their probabilities with changing $d_0$. Each curve corresponds to a single ecological network, with blue, green, red indicating $d_0 = 0.5$, $d_0 = 1.0$ and $d_0 = 2.0$, respectively. In panel **A**, there is a consistent trend of postponing saturation as $d_0$ increases, although the limit $\mathbb{P}^I$ remains unchanged. In panel **B**, there is a similar trend of postponing for achieving optimal $\mathbb{P}^M$, but the overall probability decreases with increasing $d_0$.

#### Inter-species energy exchange

The inter-species energy exchange ($\boldsymbol{\sigma}$) measures the mass-specific energy transferred from population $i$ to population $j$. In the main text, we generate these energy fluxes with a given scale $s_\sigma = 1.0$. Here, we fix the relative energy exchange network, but increase their overall scale by changing $s_\sigma \in \{1.0, 2.0, 3.0\}$ (i.e. $\boldsymbol{\sigma}' \leftarrow s_\sigma \boldsymbol{\sigma}$) to investigate how the overall scale of $\boldsymbol{\sigma}$ affects the probabilities. As seen in Fig. S4A, increase the scale of energy exchange $s_\sigma$ has no effect on initialization, consistent with previous results on feasibility analysis [15]. However, increasing $s_\sigma$ lead to a higher probability of maturation, along with a slightly larger optimal total energy supply ($Q_{\text{opt}}$).

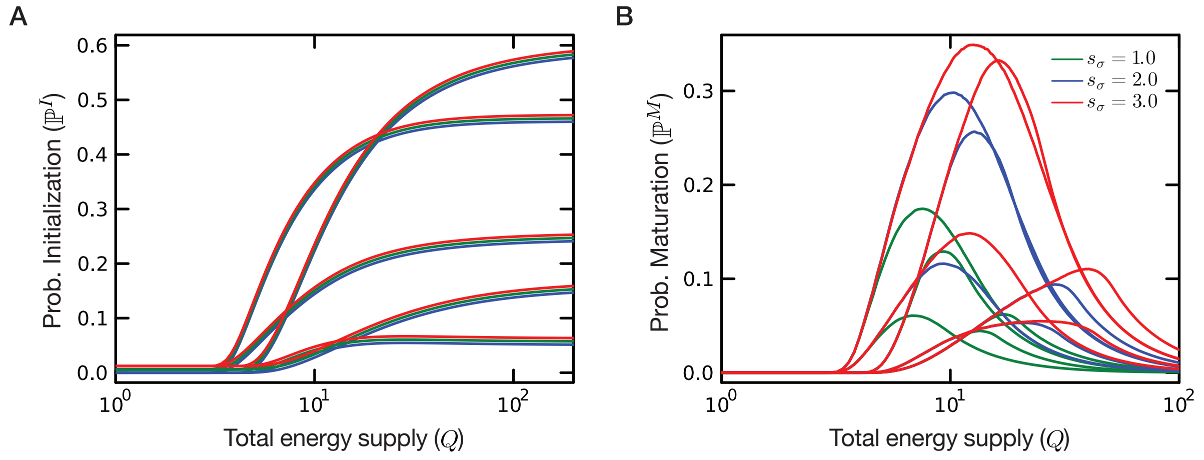


Figure S4: **Increasing scale of energy exchange has no effect on initialization, but increases the probability of maturation** Under $S = 4$, $N_0 = 1.0$, $d_0 = 1.0$, we generate five $\boldsymbol{\sigma}$ matrices and compute their probabilities (green curves), then scale these $\boldsymbol{\sigma}' \leftarrow s_\sigma \boldsymbol{\sigma}$ with $s_\sigma = 2.0$ (blue) and $s_\sigma = 3.0$ (red). **A** Probability of initialization is the same after this rescale. (Jittering to differentiate groups) **B** Probability of maturation increases as $s_\sigma$ increases, along with the slight increase of $Q_{\text{opt}}$.

#### Sensitivity to diagonal self-regulation

The dissipative ensemble used in the main text is obtained by adding a positive diagonal shift to each randomly sampled interaction matrix. Because this shift modifies only the diagonal entries of $\boldsymbol{\sigma}$, it adjusts intra-specific energy cost (self-regulation) while leaving the off-diagonal exchange structure unchanged. Here we vary the shift parameter $\sigma_0$ to test whether the main feasibility patterns depend on this choice. Such matrices are generated by drawing from random distribution and then modifying the diagonal:

> To generate such dissipative matrices, we sample $\boldsymbol{\sigma}$ from some random matrix distribution (e.g. Gaussian $\mathtt{N}(0,1)$), and we add to its diagonal elements a positive shift to guarantee the requirement of dissipative. Specifically, for any $\boldsymbol{\sigma}$, let $\sigma_0 > 0$ and $c = -\min\{\text{eigvals}(\frac{\boldsymbol{\sigma} + \boldsymbol{\sigma}^T}{2}), 0\} + \sigma_0$. Consider
>
> $$\boldsymbol{P}(c) = \frac{1}{2}((\boldsymbol{\sigma} + c\boldsymbol{I}) + (\boldsymbol{\sigma} + c\boldsymbol{I})^T) = \frac{1}{2}(\boldsymbol{\sigma} + \boldsymbol{\sigma}^T) + c\boldsymbol{I}, \tag{18}$$
>
> since $c\boldsymbol{I}$ and $\frac{1}{2}(\boldsymbol{\sigma} + \boldsymbol{\sigma}^T)$ are commutative,
>
> $$\min\{\text{eigvals}(\boldsymbol{P}(c)), 0\} = c + \min\{\text{eigvals}(\frac{1}{2}(\boldsymbol{\sigma} + \boldsymbol{\sigma}^T), 0\} = \sigma_0 > 0. \tag{19}$$
>
> Therefore, adding such $c$ to $\boldsymbol{\sigma}_{ii}$ ensures a dissipative $\boldsymbol{\sigma} + c\boldsymbol{I}$. We have chosen $\sigma_0 = 0.5$ throughout this manuscript.

Numerically, we find that an increased $\sigma_0$ slightly delays the optimal total energy supply $Q_{\text{opt}}$ in probability of maturation ($\mathbb{P}^M$) and increases the maximal probability across most $Q$ range. For initialization ($\mathbb{P}^I$), there are similar but more subtle observations. From an ecological perspective, adding $c\boldsymbol{I}$ to the energy exchange only enhances the intra-species energy cost, while the inter-species energy exchange remains unchanged. The observation that overall probabilities increase is highly consistent with previous studies on the stabilizing effects of self-regulation on ecological networks [16]. Crucially, these changes are quantitative rather than qualitative: the saturation of $\mathbb{P}^I$ and the unimodal response of $\mathbb{P}^M$ are preserved across the tested range of $\sigma_0$.

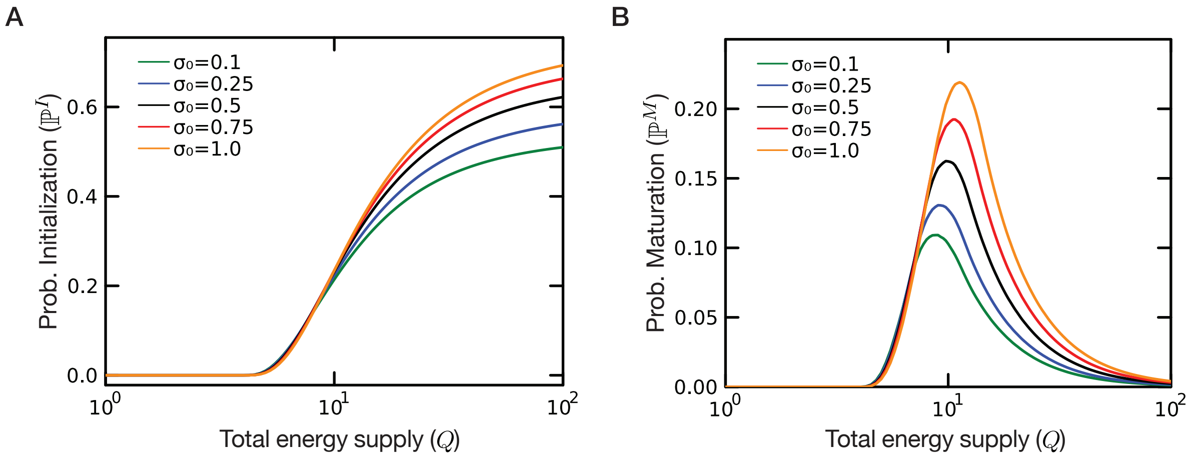


Figure S5: **Increasing intra-specific energy cost (self-regulation) enhances probabilities of initialization and maturation.** For a $S = 4$ ecological network with $N_0 = 1, d_0 = 1$, we generate one $\boldsymbol{\sigma}$ matrix and increase its diagonal part $\sigma_{ii}$ with $\sigma_0 \in [0.1, 1.0]$. Under a larger $\sigma_0$, both **(A)** the probability of initialization and **(B)** the probability of maturation increases overall. For the latter, the optimal $Q_{\text{opt}}$ delays. These trends are consistently observed for other choices of $\boldsymbol{\sigma}$.

#### Invariant scaling of energy fluxes

In the preceding sensitivity analysis, we change separately the energy exchange and utilization structures (i.e. part of $\boldsymbol{\sigma}$, $\boldsymbol{d}$). However, we also note an invariant scaling of the entire structure. As shown in this analysis, if each component of this structure is rescaled (i.e. $\boldsymbol{d} \to k\boldsymbol{d}$, $\sigma \to k\boldsymbol{\sigma}$), then the ecological network requires proportionally more energy (i.e. $Q \to kQ$) to keep up with the same probability of initialization and maturation. This is equivalent to the fact that the framework is independent of the unit to measure energy, as $k$ can be understood as a conversion factor between units.

*Statement* 8. Probability of initialization and maturation is energy unit invariant.

*Proof.* we consider $\boldsymbol{s} \in D^M(k\boldsymbol{\sigma}, k\boldsymbol{d}|kQ, \boldsymbol{N}^0)$ and $\boldsymbol{u} = k^{-1}\boldsymbol{s}$, then

$$\begin{cases} s_i \geq 0 \\ \sum_i s_i N_i^0 \leq kQ \\ \sum_i s_i (k^{-1}\boldsymbol{\sigma}^{-1}(\boldsymbol{s} - k\boldsymbol{d}))_i \leq kQ \\ (k^{-1}\boldsymbol{\sigma}^{-1}(\boldsymbol{s} - k\boldsymbol{d}))_i > 0 \end{cases} \Leftrightarrow \begin{cases} ku_i \geq 0 \\ \sum_i ku_i N_i^0 \leq kQ \\ \sum_i ku_i (k^{-1}\boldsymbol{\sigma}^{-1}(k\boldsymbol{u} - k\boldsymbol{d}))_i \leq kQ \\ (k^{-1}\boldsymbol{\sigma}^{-1}(k\boldsymbol{u} - k\boldsymbol{d}))_i > 0 \end{cases}, \tag{20}$$

which is equivalent to $\boldsymbol{u} \in D^M(\boldsymbol{\sigma}, \boldsymbol{d}|Q, \boldsymbol{N}^0)$. Therefore, $\boldsymbol{u} = k^{-1}\boldsymbol{s}$ establishes an isotropic transformation between these two maturation domains, and

$$\text{vol}(D^M(k\boldsymbol{\sigma}, k\boldsymbol{d}|kQ, \boldsymbol{N}^0)) = k^S \cdot \text{vol}(D^M(\boldsymbol{\sigma}, \boldsymbol{d}|Q, \boldsymbol{N}^0)). \tag{21}$$

Since the scaling relationship applied to each constraint in Eq. (20), the same volume scaling applied to the capture domain $D^C$ and the initialization domain $D^I$. Therefore, the $k^S$ factors appear in both sides of the probability calculations, and the probabilities remains unchanged. ■

In Fig. S6, we validate this observation numerically.

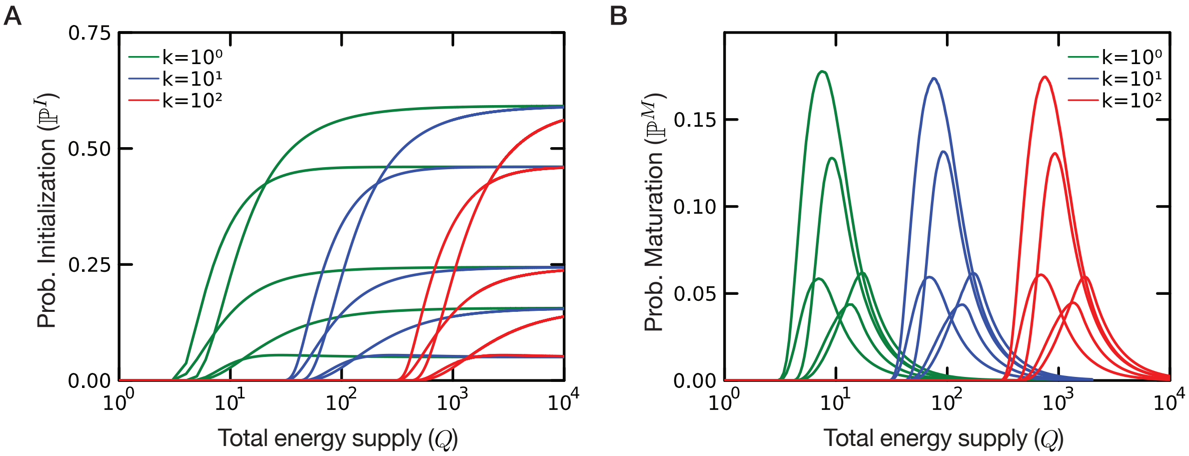


Figure S6: **Scaling $\boldsymbol{d}$ and $\boldsymbol{\sigma}$ requires a proportional $Q$ to keep the same probabilities of initialization and maturation**. For $S = 4$ ecological network with $N_0 = 1.0, d_0 = 1.0$, we generate five $\boldsymbol{\sigma}$ matrices and compute their probabilities of initialization and maturation $\mathbb{P}^R(Q)$ responses. In panels **A,B**, each green curve corresponds to the result of a single $\boldsymbol{d}, \boldsymbol{\sigma}$ (a single ecological network). We then multiply a factor of $k = 10$ (blue) and $k = 100$ (red) to these $\boldsymbol{d}$ and $\boldsymbol{\sigma}$. The new $\mathbb{P}^{I,R}(Q)$ responses are identical to the original ones up to a horizontal scale of $k$, which means a $Q' = kQ$ could recover the same level of initialization and maturation in the scaled ecological networ.

#### Minimal biomass

The minimal biomass $\boldsymbol{N}^0$ represents a fundamental constraint: $\boldsymbol{s} \cdot \boldsymbol{N}^0 \leq Q$, in the definition of all the feasibility domains. In fact, $\boldsymbol{N}^0$ determines an intrinsic scale of biomass that translates between total energy supply $Q$ and energy captures $\boldsymbol{s}$.

In the main text, we set an average minimal biomass as $N_0 = 1.0$. In this section, we relax this assumption by increasing the average level $N_0$. Numerically, we find that increasing $N_0$ does not change the limiting probability of initialization when total supply is sufficient, but delays the increase and saturation of $\mathbb{P}^I(Q)$; increasing $N_0$ consistently delays the optimal energy upper bound $Q_{\text{opt}}$, while also enhances the probability of maturation $\mathbb{P}^M(Q)$.

#### Threshold of steady-state biomass

In the main text, we always set the feasible steady state biomass condition as $\boldsymbol{N}^* > 0$; generally, we could set a threshold of feasible biomass $\boldsymbol{\epsilon} > \boldsymbol{0}$, which strengthens the condition as $N_i^* \geq \epsilon_i > 0$. Introducing such threshold generalizes our theoretical framework to broader scenarios, in particular, the partially supplied ecological network that we discussed in Section S6. There, we show that considering a partially supplied ecological network effectively raises the feasible biomass condition of autotrophs to a nonzero threshold. (refer to Eq. (25)).

Here, we perform sensitivity analysis for two cases of $\boldsymbol{\epsilon}$: 1) $\epsilon_i = 10^{-3}$ (small), 2) $\epsilon_i = 10^{-1}$ (large). We find that for small threshold level, both probabilities have negligible changes. This confirms that numerical treatment of $N^* > 0$ as $N^* \geq \epsilon$ where $\epsilon$ is a small quantity will not alter the results. For large threshold level, a higher amount of total energy supply is required for the saturation of probability of initialization, but its maximal limiting probability is unchanged; but the maximal probability of maturation is decreased.

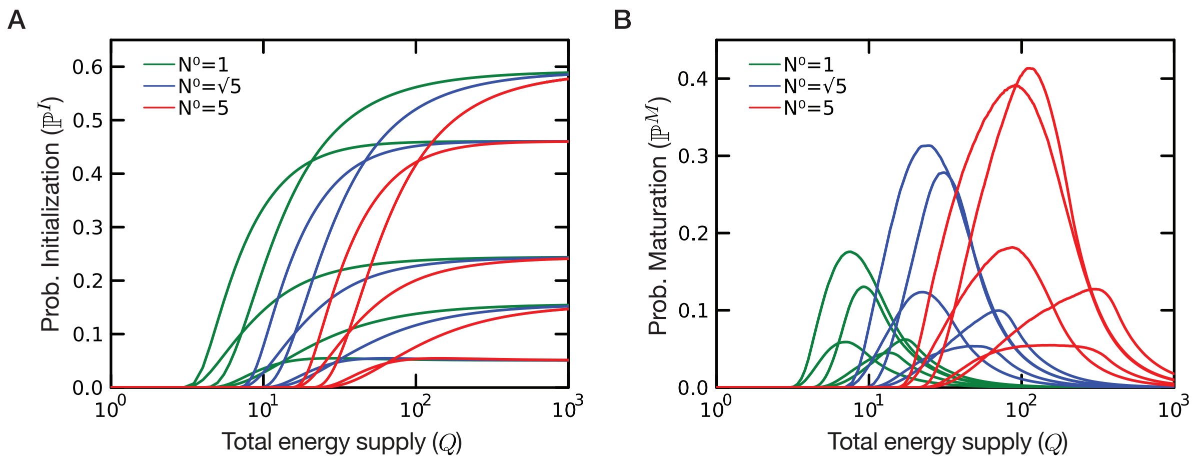


Figure S7: **Increasing minimal biomass requires larger $Q$ for initialization and maturation.** For a $S = 4$ ecological network with $d_0 = 1.0$, we generate five $\boldsymbol{\sigma}$ matrices (same as previous analysis) and increase the average minimal biomass $N_0$ from 1.0 to $\sqrt{5}$ and 5. Each green line indicates the result for a single $\boldsymbol{\sigma}$ from the original setup, while each blue (resp. red) line shows the comparison with $N_0 = \sqrt{5}$ (resp. $N_0 = 5$). Under larger $N_0$, **A** the initialization probability always saturates to the same level when $Q$ is sufficiently high, but the saturation gets delayed; **B** the maturation probability increases overall, with the optimal total energy supply $Q_{\text{opt}}$ also delayed.

#### Connectance of energy exchange

The connectance $c$ measures the probability of being nonzero for pairwise energy exchanges among the $S(S-1)$ possible off-diagonal entries of $\boldsymbol{\sigma}$. In the main text, we assume a fully connected exchange network ($c = 1.0$). Here, we vary $c \in \{1.0, 0.8, 0.6, 0.4, 0.2\}$ to investigate how network sparsity affects the probability of maturation. For each value of $c$, we generate 50 independent ecosystems (each with $S = 8$ species) by sampling distinct random seeds, and the nonzero entries of $\boldsymbol{\sigma}$ are drawn from the same distribution as in the main text. We apply functional data analysis (FDA) to each category of connetance, yielding an average trend of $P_M(Q)$; the ensemble mean and one standard deviation band are computed across the 50 replicates in logit space.

As seen in Fig. S9, four observations emerge. First, the unimodal relationship between $P_M$ and $Q$ is preserved across all connectance levels, confirming that this qualitative pattern is robust to network sparsity. Second, as connectance decreases, the optimal energy supply $Q_{\text{opt}}$ shifts systematically toward lower values, consistent with reduced internal energy exchange requiring less total supply to reach peak feasibility. Third, the median peak $P_M$ varies only modestly across connectance levels. Fourth, the variance across network realizations increases substantially at low connectance, indicating that the feasibility of sparse networks is more sensitive to the specific topology of energy exchange.

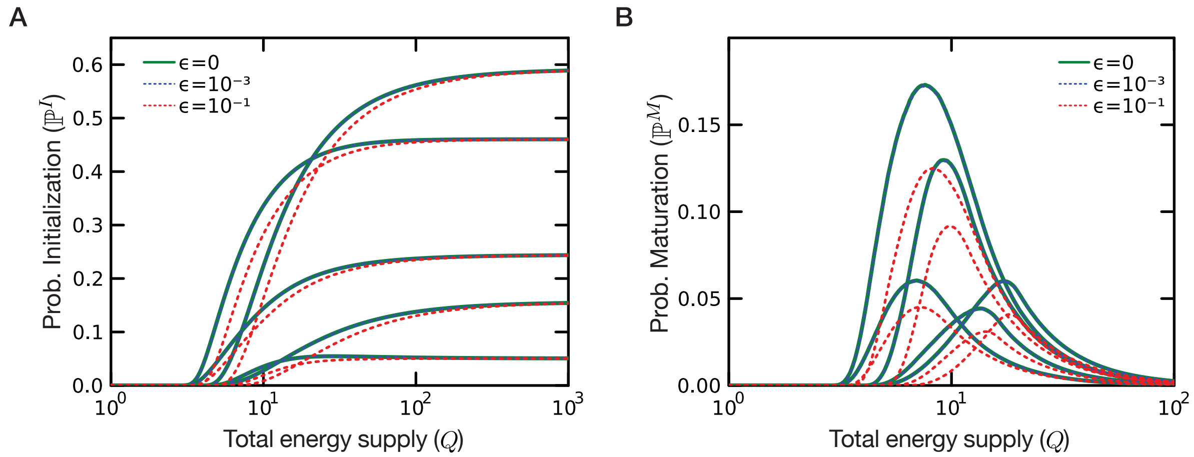


Figure S8: **Small threshold has negligible effects, while large threshold decreases probability of maturation.** For a $S = 4$ ecological network with $N_0 = 1.0$ and $d = 1.0$, we generate five $\boldsymbol{\sigma}$ matrices (same as previous analysis) and increase the threshold of steady state biomass $\epsilon_i$ from 0 to $10^{-3}$ and $10^{-1}$. Each green line indicates the result for a single $\boldsymbol{\sigma}$, while each dot blue (resp. red) line shows the comparison with $\epsilon_i = 10^{-3}$ (resp. $\epsilon_i = 10^{-1}$). **A** There is a delayed increase in initialization probability, but the limit of $\mathbb{P}^I(Q \to \infty)$ remains unchanged. **B** There is no change to the optimal $Q_{\text{opt}}$ where maturation probability is maximized; but the overall maturation probability decreases when $\epsilon_i$ increases.

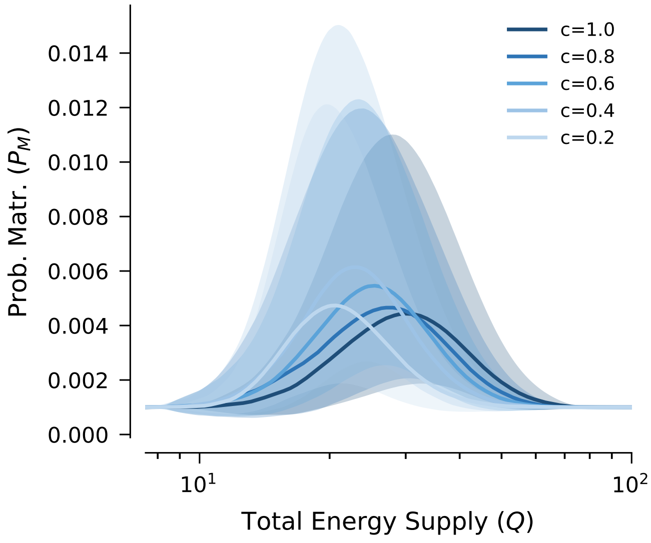


Figure S9: **Robustness of the unimodal maturation pattern to network sparsity.** Probability of maturation $P_M$ as a function of total energy supply $Q$ for networks with varying connectance $c \in \{1.0, 0.8, 0.6, 0.4, 0.2\}$, each based on 50 independent realizations of $S = 8$ networks. Solid lines show the functional median and shaded bands indicate the 68% confidence interval, both computed in logit space. The unimodal pattern is preserved at all connectance levels. Lower connectance shifts $Q_{\text{opt}}$ toward lower energy supply and increases the variability across realizations, while the median peak $P_M$ remains comparable.

### S5 The critical and optimal energy supply

In this section, we try to give an estimate of the critical and optimal energy supply. Such estimations help us to better understand the multi-phase patterns in the probability supply relationship, and provides empirical suggestions when selecting the range of $Q$ values for computation.

#### Critical energy supply

We notice that the critical energy supply that leads to a nonzero probability is *at most* the energy supply that satisfies the energy demands of a minimal biomass ecological network. We notice for most of our theoretical cases, this bound becomes an equality.

*Statement* 9. $Q_c \leq \boldsymbol{d}^\top \boldsymbol{N}^0 \equiv Q_0$

*Proof.* Consider some $Q = \boldsymbol{d}^\top \boldsymbol{N}^0 + \delta Q$ where $\delta Q > 0$, i.e., slightly higher than the bound. Then we characterize which capture $\boldsymbol{s} = \boldsymbol{d} + \delta \boldsymbol{s}$ could be feasible in the maturation domain (which is also feasible for the initialization domain). Specifically,

$$\begin{cases} d_i + \delta s_i \geq 0 \\ \sum_j (\boldsymbol{\sigma}^{-1})_{ij}\, \delta s_j > 0 \\ \sum_j \delta s_j N_j^0 \leq \delta Q \\ \sum_i \sum_j (d_i + \delta s_i)(\boldsymbol{\sigma}^{-1})_{ij} \delta s_j \leq Q \end{cases} \tag{22}$$

We see that for a sufficiently small $\delta \boldsymbol{s}$, the first and last condition can be omitted; for any given $\delta Q$, $\delta \boldsymbol{s}$ can be constructed from the traditional feasibility space of $\boldsymbol{\sigma}$, and then rescaled by the third condition.

Eventually, such $\boldsymbol{d} + \delta \boldsymbol{s}$ is feasible under the corresponding $\delta Q$, therefore the probability is nonzero. Since $\delta Q$ can be arbitrarily small, we conclude $Q_C \leq \boldsymbol{d}^\top \boldsymbol{N}^0$. ■

#### Optimal energy supply

For the unimodal pattern in $\mathbb{P}^M(Q)$, we provide a semi-quantitative analysis to estimate the optimal total energy supply ($Q_{\text{opt}}$). The overall idea is to find the condition for $Q$, at which the steady-state capture condition is the only and complete constraint on the maturation domain. Then at this point, the probability of maturation is already decreasing with larger $Q$, providing an upper bound to $Q_{\text{opt}}$.

We consider the two constraints that change with total energy supply, namely, $\boldsymbol{s}^\top \boldsymbol{N}^0 \leq Q$ $(C1)$ and $\boldsymbol{s}^\top \boldsymbol{\sigma}^{-1}(\boldsymbol{s}-\boldsymbol{d}) \leq Q$ $(C2)$. They can be linear transformed into the following standard form via $\boldsymbol{y} = \boldsymbol{L}\boldsymbol{s}$ (see S1.3, S2.3)

$$\boldsymbol{N}^{0^T}\boldsymbol{L}^{-1}\boldsymbol{y} \leq Q \;\; \text{(C1)}, \quad \boldsymbol{y}^T\boldsymbol{y} + \boldsymbol{c}^T\boldsymbol{L}^{-1}\boldsymbol{y} \leq Q \;\; \text{(C2)} \tag{23}$$

Under a given $Q$, $C1$ constrains vol$(D^C)$ (capture domain), while both C1 and C2 constrain the volume of $D^M$ (maturation domain). C1 defines a hyperplane P1: $\boldsymbol{N}^{0^T}\boldsymbol{L}^{-1}\boldsymbol{y} - Q = 0$ and C2 defines a ball B2: $||\boldsymbol{y} - \boldsymbol{y}_c|| \leq \sqrt{t}$. As $Q$ increases, the hyperplane grows faster (linear with $Q$) than the ball (sublinear with $Q$), and eventually detaches from the ball. At this point, the relative volume vol$(D^M)$/vol$(D^C)$ is already decreasing, providing an upper bound to the optimal $Q_{\text{opt}}$.

Following straightforward calculations, we can determine this point by comparing the distance from $\boldsymbol{y}_c$ to P1 and the radius of B2:

$$d(Q) := d(\boldsymbol{y}_c, P1) = \frac{|(\boldsymbol{L}^T)^{-1}\boldsymbol{N}^0 \cdot \boldsymbol{y}_c - Q|}{\|(\boldsymbol{L}^T)^{-1}\boldsymbol{N}^0\|}, \quad r(Q) := \sqrt{t}(B2) = \sqrt{Q + \frac{1}{4}\boldsymbol{c}^T\boldsymbol{P}^{-1}\boldsymbol{c}}, \tag{24}$$

which are functions of $Q$. Denote $\boldsymbol{n}^0 = \dfrac{(\boldsymbol{L}^T)^{-1}\boldsymbol{N}^0}{\|(\boldsymbol{L}^T)^{-1}\boldsymbol{N}^0\|}$ as the normal vector of P1, in the limit of $Q = 0$, $d(0) = |\boldsymbol{n}^0 \cdot \boldsymbol{y}_c| \leq \|\boldsymbol{y}_c\| = r(0)$. In the limit of $Q \to \infty$, $d(Q) \sim Q$, $r(Q) \sim Q^{\frac{1}{2}}$. Therefore, there will be an intermediate point $Q_1$ such that $d(Q_1) = r(Q_1)$, which is the upper bound for $Q_{\text{opt}}$.

#### Computational range for total energy supply

We can leverage the above bounds to select the sampling points of $Q$ at which we calculate the probability. Since computational resources is limited, we could allocate a more dense sampling around the optimal energy supply, while a loose sampling before the critical energy supply. In practice, we allocate 25 points within $[0, Q_0]$, 350 points within $[Q_0, Q_1]$ and 125 points within $[Q_1, 10Q_1]$. These $Q$ values are uniformly ranged in the logarithmic scale, which provides better numerical performance for the multi-phase Monte Carlo volume estimation (see section S2.7 and the `select_range` function).

#### S6 Partially supplied ecological networks

In real terrestrial and aquatic ecological trophic chains, only autotrophs (plants or phytoplanktons) can directly utilize free energy from the abiotic environment (i.e., $s_i > 0$ ), whereas heterotrophs cannot acquire free energy directly from the environment, but only through predation on autotrophs (i.e., $s_i = 0$). For instance, in an ecological network with 4 plants and 4 herbivores (in total $S = 8$ species), the capture domain $D_C$ should be defined as $D_C(Q, \boldsymbol{N}^0) = \{s_{1-4} > 0;\ s_{5-8} = 0;\ \sum_{i=1}^{8} s_i N_i^0 \leq Q\}$, degenerating to a 4-dimensional domain.

In the main text, we only discussed the case where all the species have $s_i \geq 0$. Then, how to study the feasibility of such *partially supplied* ecological networks? To do so, we examine each of the feasibility conditions and modify them in the new setup.

We use the subscript $a$ and $h$ to denote autotroph and heterotroph species, respectively. For the above example, $a = \{1, 2, 3, 4\}$ and $h = \{5, 6, 7, 8\}$. Without loss of generality, we assume that each index in $a$ is smaller than each index in $h$.

1) *Nonnegative energy capture:* Apparently and directly, we modify this condition to $\boldsymbol{s}_a > \boldsymbol{0}$ and $\boldsymbol{s}_h = \boldsymbol{0}$.

2) *Feasible steady state biomass:* To proceed, we denote $\boldsymbol{\sigma}^{-1} = \begin{pmatrix} \Lambda_a & \Lambda_h \end{pmatrix}$, where $\Lambda_a$ and $\Lambda_h$ is the block element of $\boldsymbol{\sigma}^{-1}$; Similarly, $\boldsymbol{s} = \begin{pmatrix} \boldsymbol{s}_a \\ \boldsymbol{0} \end{pmatrix}$ and $\boldsymbol{d} = \begin{pmatrix} \boldsymbol{d}_a \\ \boldsymbol{d}_h \end{pmatrix}$ for $\boldsymbol{s}$ and $\boldsymbol{d}$. Then, the condition reads

$$\boldsymbol{N}^* = \begin{pmatrix} \Lambda_a & \Lambda_h \end{pmatrix} \begin{pmatrix} \boldsymbol{s}_a - \boldsymbol{d}_a \\ -\boldsymbol{d}_h \end{pmatrix} > \boldsymbol{0}, \tag{25}$$

which can be simplified as $\boldsymbol{N}_a^* = \Lambda_a(\boldsymbol{s}_a - \boldsymbol{d}_a) > \Lambda_h \boldsymbol{d}_h$. Notice that the lower bound for the steady state biomass of autotrophs is increased from $\boldsymbol{0}$ to a non-zero value $\Lambda_h \boldsymbol{d}_h$, which is associated with the demands of heterotrophs $\boldsymbol{d}_h$.

3) *Minimal energy capture.* Since $\boldsymbol{s}_h = \boldsymbol{0}$, the condition is reduced to $\sum_{i \in a} s_i N_i^0 \leq Q$.

4) *Feasible steady-state energy capture* Since $\boldsymbol{s}_h = \boldsymbol{0}$, the condition is reduced to $\sum_{i \in a} s_i N_i^* \leq Q$.

Surprisingly, we notice a simple mapping from the feasibility domain of the partially supplied network to that of an fully supplied network of all the autotrophs, with only a reduction of dimensions and an increase of the threshold for steady-state biomass. Immediately, this suggests our computational framework is applicable to such general case, and the qualitative patterns remain similar (see S4.6).

### S7 Density-dependent capture rates

In the main text, we have followed a per-capita form to calculate the flow of energies within the community and between environment-community. Specifically, if the biomass population $i$ is $N_i$, then the total energy captured by the network at unit time is $Q(\boldsymbol{s}, \boldsymbol{N}) = \sum_{i=1}^{S} s_i N_i$. Specifically, at equilibrium, total energy capture is $Q(\boldsymbol{s}, \boldsymbol{N}^*(\boldsymbol{s})) = \sum_{i=1}^{S} s_i \sum_{j=1}^{S} (\sigma^{-1})_{ij}(s_j - d_j)$. While this key assumption has facilitated the derivation and computation of feasibility domains, it also imposes a global linear relationship in $\boldsymbol{N}^*(\boldsymbol{s}) = \boldsymbol{\sigma}^{-1}(\boldsymbol{s} - \boldsymbol{d})$. However, real-world ecological dynamics are often governed by nonlinear interactions and saturating uptake kinetics that prevent such "unbounded response"; to ensure the robustness of our result on the probability of maturation ($\mathbb{P}_M$), we here extend the framework to incorporate density-dependent capture rates that mimic the realistic negative feedback between population size and per-capita capture rates.

We consider the following hybrid dynamics (26), where the energy capture rate term is density-dependent with a decaying coefficient $\theta(\vec{N})$. Such term can be viewed as resource-mediated, which introduces a natural saturating effect on the per-capita capture rates with $\dfrac{\partial \theta(\vec{N})}{\partial N_j} < 0$. At the same time, we still keep the direct flows energy exchange term $\sigma_{ij}$.

$$\frac{\mathrm{d}N_i}{\mathrm{d}t} = N_i \left( s_i \theta(\vec{N}) - d_i - \sum_j \sigma_{ij} N_j \right) \tag{26}$$

In this study, we consider a linear perturbation form of $\theta(\vec{N}) = 1 - \sum_j k_j N_j, k_j > 0$. Under such assumption, (26) still follows the generalized Lotka-Volterra form, but the interaction matrix is perturbed from $\sigma$ by adding a $sk^T$ term. The larger $\|k\|$, the stronger such perturbation.

**The equilibrium biomass and feasibility constraint for** (26)

The equilibrium biomass is determined as

$$\boldsymbol{N}^*(\boldsymbol{k}) = \underbrace{(\boldsymbol{\sigma} + \boldsymbol{s}k^\top)^{-1}}_{\triangleq \Lambda(k)}(\boldsymbol{s} - \boldsymbol{d}), \tag{27}$$

by using the Sherman–Morrison formula, we have

$$\Lambda(k) = \boldsymbol{\sigma}^{-1} - \boldsymbol{\sigma}^{-1} \frac{\boldsymbol{s}k^\top}{1 + k^\top \boldsymbol{\sigma}^{-1}\boldsymbol{s}} \boldsymbol{\sigma}^{-1}. \tag{28}$$

Let us denote $\boldsymbol{N}^* = \boldsymbol{\sigma}^{-1}(\boldsymbol{s} - \boldsymbol{d})$ the unperturbed equilibrium, $\alpha = 1 + \boldsymbol{k}^\top \boldsymbol{\sigma}^{-1} \boldsymbol{d}$ as a constant scalar.

After some algebra, it can be shown that

$$\boldsymbol{N}^*(\boldsymbol{k}) = \frac{(\alpha \boldsymbol{I} - \boldsymbol{\sigma}^{-1}\boldsymbol{d}\boldsymbol{k}^\top)}{\alpha + \boldsymbol{k}^\top \boldsymbol{N}^*}\boldsymbol{N}^* = \frac{1}{\alpha + \boldsymbol{k}^\top \boldsymbol{N}^*}\boldsymbol{P}(\boldsymbol{k})(\boldsymbol{s} - \boldsymbol{d}), \tag{29}$$

where $\boldsymbol{P}(\boldsymbol{k}) = \alpha\boldsymbol{\sigma}^{-1} - \boldsymbol{\sigma}^{-1}\boldsymbol{d}\boldsymbol{k}^\top\boldsymbol{\sigma}^{-1}$.

Notice that both the denominator and the numerator is linearly dependent of $\boldsymbol{s}$, which leads to a nonlinear, saturating relationship between $\boldsymbol{N}^*(\boldsymbol{k})$ and $\boldsymbol{s}$. In this line, the **feasible equilibrium biomass** condition becomes

$$\boldsymbol{P}(\boldsymbol{k})(\boldsymbol{s} - \boldsymbol{d}) > \boldsymbol{0} \tag{30}$$

The maturation energy capture condition is $\sum_i s_i N^*(\boldsymbol{k})_i \leq Q$. Under some algebra, this condition is simplified as

$$\boldsymbol{s}^\top \boldsymbol{P}(\boldsymbol{k})\boldsymbol{s} - [\boldsymbol{P}(\boldsymbol{k})\boldsymbol{d} + Q(\boldsymbol{\sigma}^{-1})^\top \boldsymbol{k}]^\top \boldsymbol{s} \leq Q \tag{31}$$

Surprisingly, both (30) and (31) follow the same form as in S2.3, which grants us the same methods to be applied to sample and compute the modified feasibility domains.

In Figure S10, we randomly sampled and computed the probability of maturation for 25 model networks under increasing $k$. We find that increasing the average saturation strength $k$ creates a continuous spectrum of relationships between the probability of maturation ($\mathbb{P}_M$) and total energy supply ($Q$). First, when $k \to 0$, the model recovers our primary conclusion, where $\mathbb{P}_M$ follows a strong unimodal response that eventually decays to zero at high $Q$. Second, within a range of saturation strength ($k > 0$), the unimodal response is preserved but shifts quantitatively: the optimal energy supply ($Q_{\text{opt}}$) increases across scales, but the decay of $\mathbb{P}_M$ at high $Q$ is buffered. The stronger saturation (large $k$), the response transitions from a sharp peak to a scattered one. Crucially, unlike the the previous linear case, $\mathbb{P}_M$ converges to a non-zero value for large $Q$. For larger $S$ and high $k$, the response can degenerate to a saturating shape when this limiting probability is high.

This transition reveals the fundamental interplay between energetic constraints and ecological feasibility. In the low-supply regime, the saturation effect is negligible since $s_i k_j$ is small, therefore the feasibility of networks is identical to the unperturbed, linear model. As $Q$ increases, however, the saturation term acts as self-regulatory mechanism: while the linear model predicts that excessive energy supply decreases feasibility by driving biomass beyond feasible boundaries, the saturation mechanism converts excess energy into stronger competitive feedback and leads to a non-vanishing, saturated probability of maturation. Importantly, the existence of an optimal total energy supply ($Q_{\text{opt}}$) remains a robust feature within a wide range of saturation strength, confirming that the unimodal response is not an artifact of linearity but a general property of energy-constrained networks, modulated by the strength of resource saturation.

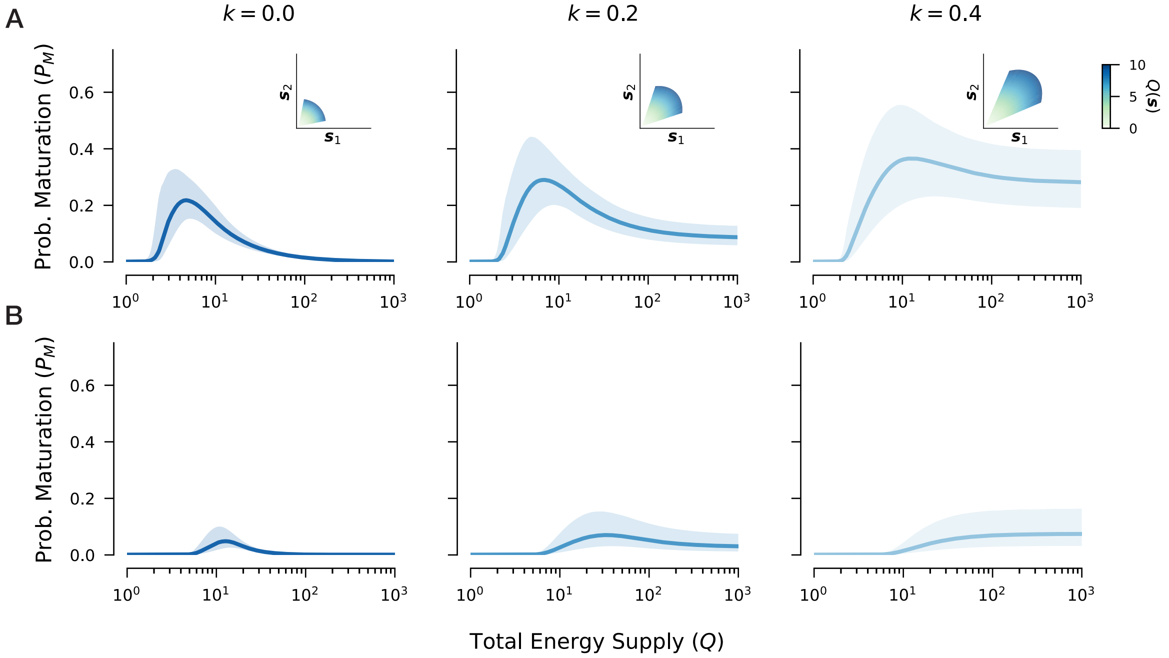


Figure S10: **Robustness of feasibility patterns under density-dependent energy capture. Main panels**: Expected probabilities of maturation ($P_M$) as a function of total energy supply ($Q \in [10^0, 10^3]$) for varying saturation strengths $k \in \{0, 0.2, 0.4\}$. Probabilities are calculated from 25 independent replicates using randomly sampled network parameters $(\boldsymbol{\sigma}, \boldsymbol{d}, \boldsymbol{N}^0)$ at **(A)** $S = 2$ and **(B)** $S = 4$. As the saturation strength $k$ increases, the unimodal response and the existence of an optimal energy supply ($Q_{\text{opt}}$) remain as robust patterns. Notably, for $k > 0$, the limiting probability at $Q \to \infty$ converges to a non-zero plateau rather than vanishing, indicating that saturation mechanisms alleviate energetic constraints at the limit of large supply. **Insets**: Geometric characterization of the maturation domain ($D_M$) and equilibrium total capture $Q(\boldsymbol{s}) = \boldsymbol{s} \cdot \boldsymbol{N}^*$ for a selected model network at $Q = 10$. The saturating strength $k$ lowers the steady-state biomass $\mathbf{N}^*$ for a given energy capture rate $\mathbf{s}$. This effect results in a slight angular shrinkage but a significant radial expansion of $D_M$. These geometric observations explain the increase in limiting probability and the persistence of the unimodal pattern across different saturation regimes.

### S8 Extension to feasibility partitions

In the main text and the previous sections, the *feasibility domain* refers to the full-coexistence case, where all $S$ populations persist simultaneously. Here we extend the same construction to all candidate communities $\mathcal{C} \subseteq \{1, \cdots, S\}$. We will use *feasibility partition* to denote the subset of $\boldsymbol{s}$-space associated with one fixed community $\mathcal{C}$, and *feasibility partitions* to denote the full collection over the $2^S$ possible communities. Under this convention, the original full-community feasibility domain is simply the partition associated with $\mathcal{C} = \{1, \cdots, S\}$.

#### Complementarity condition in feasibility partitions

For a fixed community $\mathcal{C} \subseteq \{1, \cdots, S\}$, we first define its *unconstrained feasibility partition*: the set of capture vectors $\boldsymbol{s}$ that admit a boundary equilibrium with support exactly on $\mathcal{C}$, without yet imposing any constraint on the total energy supply $Q$. A standard way to express this requirement is through complementarity between biomass and per-capita net growth. Define the per-capita net growth vector

$$\boldsymbol{g}(\boldsymbol{N}; \boldsymbol{s}) = \boldsymbol{s} - \boldsymbol{d} - \boldsymbol{\sigma} \boldsymbol{N}, \tag{32}$$

whose $i$-th component is $g_i(\boldsymbol{N}; \boldsymbol{s}) = s_i - d_i - \sum_j \sigma_{ij} N_j$. Then a boundary equilibrium with support $\mathcal{C}$ is characterized by the strict complementarity conditions

$$\begin{cases} N_i > 0, \quad g_i(\boldsymbol{N}; \boldsymbol{s}) = 0, \quad i \in \mathcal{C}, \\ N_i = 0, \quad g_i(\boldsymbol{N}; \boldsymbol{s}) < 0, \quad i \notin \mathcal{C}. \end{cases} \tag{33}$$

Equivalently, one may write $N_i \geq 0$, $g_i(\boldsymbol{N}; \boldsymbol{s}) \leq 0$, and $N_i g_i(\boldsymbol{N}; \boldsymbol{s}) = 0$ for all $i$, together with $\mathrm{supp}(\boldsymbol{N}) = \mathcal{C}$ and the strict inequality $g_i(\boldsymbol{N}; \boldsymbol{s}) < 0$ for extinct populations. The following representation makes this condition explicit [17]:

*Statement* 10. For a fixed community $\mathcal{C} \subseteq \{1, \cdots, S\}$,

$$D_f(\mathcal{C}) = \left\{ \boldsymbol{s} = \boldsymbol{d} + \sum_{i \in \mathcal{C}} n_i \boldsymbol{\sigma}_i - \sum_{i \notin \mathcal{C}} n_i \boldsymbol{e}_i \mid n_i > 0 \quad \forall 1 \leq i \leq S \right\}, \tag{34}$$

where $\boldsymbol{\sigma}_i$ is the $i$-th column of $\boldsymbol{\sigma}$, and $\boldsymbol{e}_i$ is the $i$-th unit vector, namely, $\boldsymbol{e}_i[j] = \mathbb{1}(j = i)$ for all $1 \leq j \leq S$. This set is exactly the set of capture vectors $\boldsymbol{s}$ for which there exists a biomass vector $\boldsymbol{N}$ satisfying the strict complementarity conditions in Eq. (33).

*Proof.* Suppose first that $\boldsymbol{s} \in D_f(\mathcal{C})$. Then there exist positive numbers $\{n_i\}_{i=1}^S$ such that

$$\boldsymbol{s} = \boldsymbol{d} + \sum_{i \in \mathcal{C}} n_i \boldsymbol{\sigma}_i - \sum_{i \notin \mathcal{C}} n_i \boldsymbol{e}_i. \tag{35}$$

Define a biomass vector $\boldsymbol{N}$ by $N_i = n_i$ for $i \in \mathcal{C}$ and $N_i = 0$ for $i \notin \mathcal{C}$. Since $\sum_{i \in \mathcal{C}} n_i \boldsymbol{\sigma}_i = \boldsymbol{\sigma N}$, we obtain

$$\boldsymbol{g}(\boldsymbol{N}; \boldsymbol{s}) = \boldsymbol{s} - \boldsymbol{d} - \boldsymbol{\sigma N} = -\sum_{i \notin \mathcal{C}} n_i \boldsymbol{e}_i. \tag{36}$$

Therefore, $g_i(\boldsymbol{N}; \boldsymbol{s}) = 0$ for all $i \in \mathcal{C}$, while $g_i(\boldsymbol{N}; \boldsymbol{s}) = -n_i < 0$ for all $i \notin \mathcal{C}$. Together with $N_i > 0$ on $\mathcal{C}$ and $N_i = 0$ on its complement, Eq. (33) follows.

Conversely, suppose there exists a biomass vector $\boldsymbol{N}$ satisfying Eq. (33). Let $n_i = N_i$ for $i \in \mathcal{C}$ and $n_i = -g_i(\boldsymbol{N}; \boldsymbol{s})$ for $i \notin \mathcal{C}$. By assumption, all these coefficients are strictly positive. Since $N_i = 0$ for $i \notin \mathcal{C}$, we have

$$\boldsymbol{\sigma N} = \sum_{i \in \mathcal{C}} N_i \boldsymbol{\sigma}_i = \sum_{i \in \mathcal{C}} n_i \boldsymbol{\sigma}_i. \tag{37}$$

Using $\boldsymbol{g}(\boldsymbol{N}; \boldsymbol{s}) = \boldsymbol{s} - \boldsymbol{d} - \boldsymbol{\sigma N}$, we then obtain

$$\boldsymbol{s} = \boldsymbol{d} + \boldsymbol{\sigma N} + \boldsymbol{g}(\boldsymbol{N}; \boldsymbol{s}) = \boldsymbol{d} + \sum_{i \in \mathcal{C}} n_i \boldsymbol{\sigma}_i - \sum_{i \notin \mathcal{C}} n_i \boldsymbol{e}_i, \tag{38}$$

which is precisely Eq. (34). Hence $\boldsymbol{s} \in D_f(\mathcal{C})$. ■

This representation is useful because it rewrites the complementarity condition entirely as a geometric constraint on $\boldsymbol{s}$, depending only on $\boldsymbol{d}$ and $\boldsymbol{\sigma}$. In particular, $D_f(\mathcal{C})$ is an unconstrained feasibility partition with linear boundaries: it is an affine image of the positive orthant generated by the columns $\{\boldsymbol{\sigma}_i\}_{i \in \mathcal{C}}$ and $\{-\boldsymbol{e}_i\}_{i \notin \mathcal{C}}$. Therefore, before introducing the energy-supply constraint $Q$, each partition already has a polyhedral structure and remains directly computable by standard linear-geometry methods.

#### Solving boundary equilibrium

Once the support $\mathcal{C}$ is fixed, the equilibrium biomass is obtained by solving only the surviving subsystem. The main technical step is then to re-embed this subsystem equilibrium into the original $S$-dimensional coordinates, so that all later energetic constraints can still be written in the same linear/quadratic form used by our unified *InterPolyBalls* framework.

Let $\mathcal{O} = \{1, \cdots, S\} \setminus \mathcal{C}$ denote the complement of $\mathcal{C}$. For any vector $\boldsymbol{x}$ and matrix $\boldsymbol{A}$, write

$$\boldsymbol{x}_{\mathcal{C}} = (x_i)_{i\in\mathcal{C}}, \qquad \boldsymbol{x}_{\mathcal{O}} = (x_i)_{i\in\mathcal{O}}, \qquad \boldsymbol{A}_{\mathcal{CC}} = (A_{ij})_{i,j\in\mathcal{C}}.$$

Choose a permutation matrix $M$ that places the indices in $\mathcal{C}$ first, so that

$$M^\top \boldsymbol{x} = \begin{pmatrix} \boldsymbol{x}_{\mathcal{C}} \\ \boldsymbol{x}_{\mathcal{O}} \end{pmatrix}, \qquad M^\top \boldsymbol{A} M = \begin{pmatrix} \boldsymbol{A}_{\mathcal{CC}} & \boldsymbol{A}_{\mathcal{CO}} \\ \boldsymbol{A}_{\mathcal{OC}} & \boldsymbol{A}_{\mathcal{OO}} \end{pmatrix}.$$

Then, for a given $\boldsymbol{s} \in D_f(\mathcal{C})$, the boundary equilibrium supported on $\mathcal{C}$ is

$$\boldsymbol{N}^*_{\mathcal{C}} = (\boldsymbol{\sigma}_{\mathcal{CC}})^{-1}(\boldsymbol{s}_{\mathcal{C}} - \boldsymbol{d}_{\mathcal{C}}), \qquad \boldsymbol{N}^*_{\mathcal{O}} = \boldsymbol{0}.$$

Equivalently, in the reordered coordinates,

$$\begin{pmatrix} \boldsymbol{N}^*_{\mathcal{C}} \\ \boldsymbol{N}^*_{\mathcal{O}} \end{pmatrix} = \begin{pmatrix} (\boldsymbol{\sigma}_{\mathcal{CC}})^{-1} & \boldsymbol{0} \\ \boldsymbol{0} & \boldsymbol{0} \end{pmatrix} \begin{pmatrix} \boldsymbol{s}_{\mathcal{C}} - \boldsymbol{d}_{\mathcal{C}} \\ \boldsymbol{s}_{\mathcal{O}} - \boldsymbol{d}_{\mathcal{O}} \end{pmatrix}, \qquad \begin{pmatrix} \boldsymbol{s}_{\mathcal{C}} - \boldsymbol{d}_{\mathcal{C}} \\ \boldsymbol{s}_{\mathcal{O}} - \boldsymbol{d}_{\mathcal{O}} \end{pmatrix} = M^\top(\boldsymbol{s} - \boldsymbol{d}).$$

Translated back to the original indices,

$$\boldsymbol{N}^* = \underbrace{M \begin{pmatrix} \boldsymbol{N}^*_{\mathcal{C}} \\ \boldsymbol{N}^*_{\mathcal{O}} \end{pmatrix}}_{\text{restore original ordering}} = \underbrace{M \begin{pmatrix} (\boldsymbol{\sigma}_{\mathcal{CC}})^{-1} & \boldsymbol{0} \\ \boldsymbol{0} & \boldsymbol{0} \end{pmatrix} M^\top}_{=\boldsymbol{\Lambda}} (\boldsymbol{s} - \boldsymbol{d}) \equiv \boldsymbol{\Lambda}(\boldsymbol{s} - \boldsymbol{d}). \tag{39}$$

Notice that $\boldsymbol{\Lambda}$ has the full dimension $S$, and therefore generalizes the role of $\boldsymbol{\sigma}^{-1}$ from the full-community feasibility domain to the community-specific feasibility partitions. With Eq. (39), we can readily translate the energetic constraints into functions of $\boldsymbol{s}$, and therefore impose geometric constraints directly on $D_f(\mathcal{C})$. In the next subsection, we provide the road map from this unconstrained partition to its maturation-stage generalization under the energy-supply constraint.

#### Energetic constraints on feasibility partition

To obtain the constrained partitions at a given supply level $Q$, we must intersect $D_f(\mathcal{C})$ with the positive-capture condition $\boldsymbol{s} > \boldsymbol{0}$ and the energetic condition $\sum_i s_i N_i(t) \leq Q$, where $Q$ is the total energy supply from the environment. As before, we focus on the initialization stage $t = 0$, where $\sum_i s_i N_i^0 \leq Q$, and the maturation stage $t \to \infty$, where $\sum_i s_i N_i^* \leq Q$. Thus, the *initialization partition* is obtained by intersecting $D_f(\mathcal{C})$ with the common constraints $\boldsymbol{s} > \boldsymbol{0}$ and $\sum_i s_i N_i^0 \leq Q$, whereas the *maturation partition* is obtained by further intersecting with a community-specific quadratic constraint induced by the corresponding boundary equilibrium. In particular,

$$D_I(\mathcal{C}; \boldsymbol{\sigma}, \boldsymbol{d} \mid Q, \boldsymbol{N}^0) = \{\boldsymbol{s} \in D_f(\mathcal{C}) \mid \boldsymbol{s} > \boldsymbol{0},\ \boldsymbol{s}^\top \boldsymbol{N}^0 \leq Q\}, \tag{40}$$

and

$$D_M(\mathcal{C}; \boldsymbol{\sigma}, \boldsymbol{d} \mid Q, \boldsymbol{N}^0) = \{\boldsymbol{s} \in D_I(\mathcal{C}; \boldsymbol{\sigma}, \boldsymbol{d} \mid Q, \boldsymbol{N}^0) \mid \boldsymbol{s} > \boldsymbol{0},\ \boldsymbol{s}^\top \boldsymbol{N}^* \leq Q\}. \tag{41}$$

*Statement* 11. The energetic constraint at maturation can be expressed as a community-specific quadratic constraint. Specifically,

$$\boldsymbol{s}^\top \boldsymbol{P}\boldsymbol{s} + \boldsymbol{h}^\top \boldsymbol{s} \le Q, \tag{42}$$

Where $P = \frac{1}{2}(\boldsymbol{\Lambda} + \boldsymbol{\Lambda}^\top), \boldsymbol{h} = -\boldsymbol{\Lambda}\boldsymbol{d}$, whose $\boldsymbol{\Lambda}$ is defined in (39).

*Proof.* For a fixed community $\mathcal{C}$, we start from the same energetic principle $\sum_i s_i N_i^* = \boldsymbol{s}^\top \boldsymbol{N}^* \le Q$. Using Eq. (39), we obtain

$$\boldsymbol{s}^\top \boldsymbol{\Lambda}(\boldsymbol{s} - \boldsymbol{d}) \le Q. \tag{43}$$

Separating the quadratic and linear terms, and symmetrizing the quadratic form, gives the above form. ■

#### Computation of constrained feasibility partitions

While the computation of initialization partition involves adding the common constraints $\boldsymbol{s} > \boldsymbol{0}$ and $\sum_i s_i N_i^0 \le Q$ on $D_f(\mathcal{C})$ and is quite straightforward to implement, the computation of maturation partition requires further generalization of current pipeline. The specific challenges lies in the rank of $\boldsymbol{P}$, which is at most $|\mathcal{C}|$. This changes the geometric nature of the maturation constraint from a full sphere to a cylinder.

*Statement* 12. Consider the following $\boldsymbol{L}_0$ as the $|\mathcal{C}| \times |\mathcal{C}|$ Cholesky matrix such that

$$\boldsymbol{L}_0^\top \boldsymbol{L}_0 = \frac{1}{2}(\boldsymbol{\sigma}_{\mathcal{CC}})^{-1} + \frac{1}{2}((\boldsymbol{\sigma}_{\mathcal{CC}})^{-1})^\top. \tag{44}$$

and construct the following linear transformation:

$$\boldsymbol{L} = \begin{pmatrix} \boldsymbol{L}_0 & \boldsymbol{0} \\ \boldsymbol{0} & \boldsymbol{I} \end{pmatrix} M^\top. \tag{45}$$

Then, the linear transformation $\boldsymbol{y} = \boldsymbol{L}\boldsymbol{s}$ maps constraint (42) into a standard form: either a sphere for the full community, or a cylinder for partial communities.

Once we construct such linear transformation, we can compute and analyze the maturation partition (constrained feasibility partition) using methods already developed in S2, where the exact implementation of sampling and volume estimation techniques remain the same. Since we are dealing with not only spheres but also cylinders, we shall refer to the standardized domains as *InterPolyQuads* in our software implementation.

Lastly, we give the proof for the above statement as well as the detailed form of constraint after the linear transform.

*Proof.* Accroding to previous definition,

$$\boldsymbol{P} = M \begin{pmatrix} \frac{1}{2}(\boldsymbol{\sigma}_{\mathcal{CC}})^{-1} + \frac{1}{2}((\boldsymbol{\sigma}_{\mathcal{CC}})^{-1})^\top & \mathbf{0} \\ \mathbf{0} & \mathbf{0} \end{pmatrix} M^\top, \tag{46}$$

which has $\text{rank}(\boldsymbol{P}) = |\mathcal{C}|$.

Let $\boldsymbol{y} = \boldsymbol{L}\boldsymbol{s}$. Under the inverse transformation $\boldsymbol{s} = \boldsymbol{L}^{-1}\boldsymbol{y}$, the quadratic term is standardized as

$$\begin{aligned} \boldsymbol{s}^\top \boldsymbol{P} \boldsymbol{s} &= \boldsymbol{y}^\top \begin{pmatrix} (\boldsymbol{L}_0^{-1})^\top & \mathbf{0} \\ \mathbf{0} & \boldsymbol{I} \end{pmatrix} M^\top \boldsymbol{P} M \begin{pmatrix} \boldsymbol{L}_0^{-1} & \mathbf{0} \\ \mathbf{0} & \boldsymbol{I} \end{pmatrix} \boldsymbol{y} \\ &= \boldsymbol{y}^\top \begin{pmatrix} (\boldsymbol{L}_0^{-1})^\top \left(\frac{1}{2}(\boldsymbol{\sigma}_{\mathcal{CC}})^{-1} + \frac{1}{2}((\boldsymbol{\sigma}_{\mathcal{CC}})^{-1})^\top\right) \boldsymbol{L}_0^{-1} & \mathbf{0} \\ \mathbf{0} & \mathbf{0} \end{pmatrix} \boldsymbol{y} \\ &= \boldsymbol{y}^\top \begin{pmatrix} \boldsymbol{I}_{|\mathcal{C}|} & \mathbf{0} \\ \mathbf{0} & \mathbf{0} \end{pmatrix} \boldsymbol{y} \\ &\equiv \boldsymbol{y}^\top \Pi \boldsymbol{y}. \end{aligned} \tag{47}$$

Where $\Pi$ is a rank-$|\mathcal{C}|$ projection matrix with ones on the first $|\mathcal{C}|$ diagonal elements and zero elsewhere.

Then, define $\boldsymbol{y}_c = -\frac{1}{2}(\boldsymbol{L}^{-1})^\top \boldsymbol{h}$,

$$\begin{aligned} \boldsymbol{y}_c = \frac{1}{2}(\boldsymbol{L}^{-1})^\top \boldsymbol{\Lambda} \boldsymbol{d} &= \frac{1}{2} \begin{pmatrix} (\boldsymbol{L}_0^{-1})^\top & \mathbf{0} \\ \mathbf{0} & \boldsymbol{I} \end{pmatrix} M^\top M \begin{pmatrix} (\boldsymbol{\sigma}_{\mathcal{CC}})^{-1} & \mathbf{0} \\ \mathbf{0} & \mathbf{0} \end{pmatrix} M^\top \boldsymbol{d} \\ &= \frac{1}{2} \begin{pmatrix} (\boldsymbol{L}_0^{-1})^\top (\boldsymbol{\sigma}_{\mathcal{CC}})^{-1} \boldsymbol{d}_{\mathcal{C}} \\ \mathbf{0} \end{pmatrix}, \end{aligned} \tag{48}$$

whose entries after row $|\mathcal{C}|$ are bound to be zero. Therefore, $\Pi \boldsymbol{y}_c \equiv \boldsymbol{y}_c$, and the constraint takes the standardized form

$$\begin{aligned} \boldsymbol{s}^\top \boldsymbol{P} \boldsymbol{s} + \boldsymbol{h}^\top \boldsymbol{s} = \boldsymbol{y}^\top \Pi \boldsymbol{y} - 2\boldsymbol{y}_c^\top \boldsymbol{y} &\leq Q \\ \Leftrightarrow (\boldsymbol{y} - \boldsymbol{y}_c)^\top \Pi (\boldsymbol{y} - \boldsymbol{y}_c) &\leq Q + \boldsymbol{y}_c^\top \boldsymbol{y}_c. \end{aligned} \tag{49}$$

We note the last quadratic inequality generally defines the region surrounded by a cylinder. ∎

Using the geometric characterization and computational procedure developed above, we can now extend the partition analysis in Fig. 4 of the main text to additional network sizes. In particular, we repeat the same calculation over independent network replicates for $S \in \{2, 4, 8\}$, so that the resulting patterns can be compared directly with the $S = 6$ case shown in the main text.

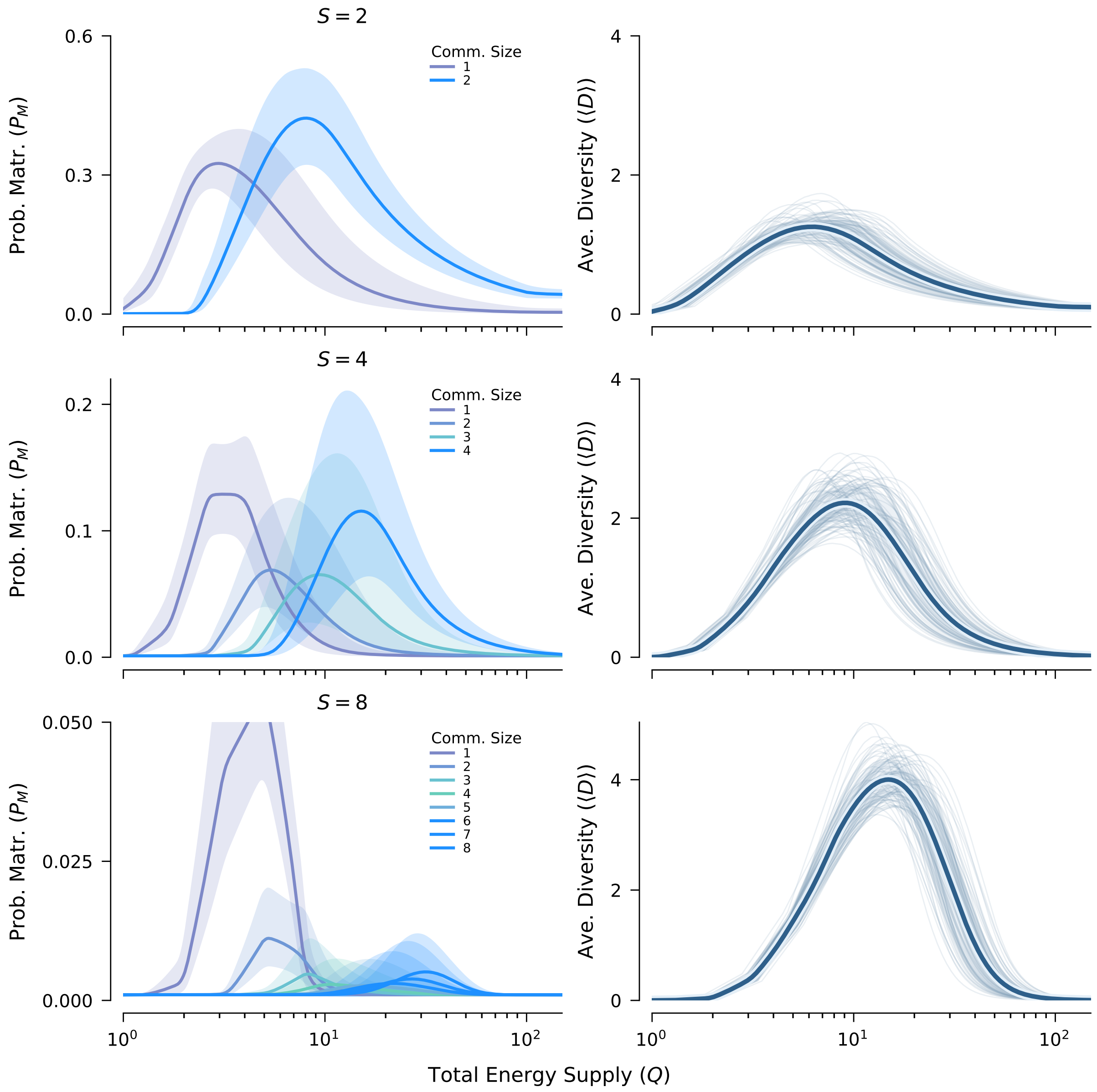


Figure S11: **Feasibility partitions across additional network sizes.** Using the computational pipeline developed above, we replicate the analysis of Fig. 4 in the main text for additional network sizes $S \in \{2, 4, 8\}$. As in Fig. 4, each row summarizes 100 independent network replicates for one fixed value of $S$. For each size, the left panel shows the probability of maturation $P_M$ as a function of total energy supply $Q$ for candidate communities grouped by community size, and the right panel shows the corresponding average diversity $\langle D \rangle$. Across sizes, the probability of maturation remains unimodal within each community-size class, and the energetic window shifts upward as community size increases. The same patterns are observed for average diversity as well.

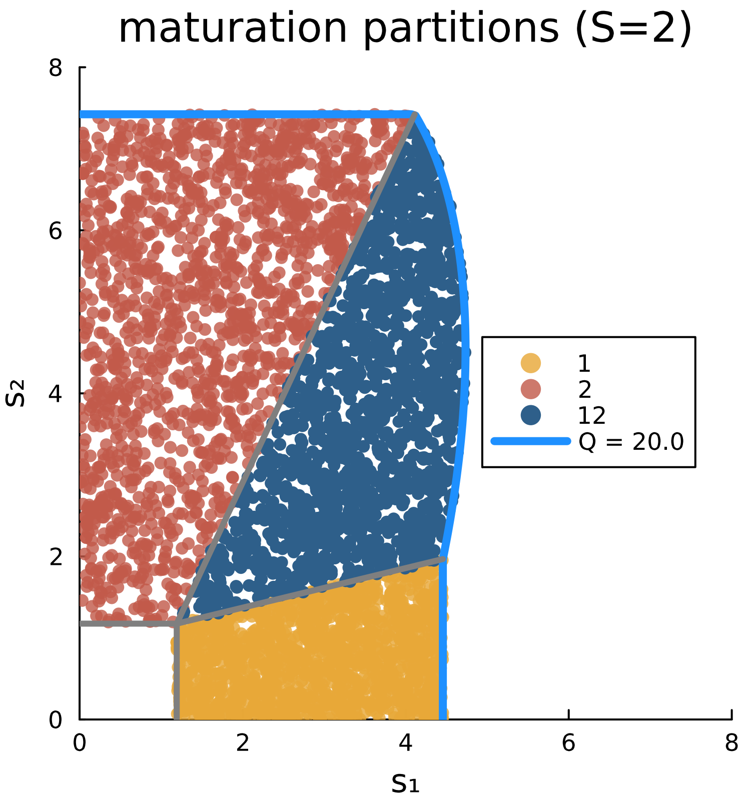


Figure S12: **Visualizing maturation partitions for a $S = 2$ model network at $Q = 20.0$.** Each colored region represents 1500 samples drawn uniformly from the maturation domain $D_M(\mathcal{C})$ of a candidate community $\mathcal{C}$: the full community $\{1, 2\}$ (dark blue), and the partial communities $\{1\}$ (orange) and $\{2\}$ (red). Gray lines indicate the boundaries of the classical feasibility domain $D_f(\mathcal{C})$, which partition $\boldsymbol{s}$-space by requiring all active populations to maintain positive biomass. The outer blue curve marks the energy constraint boundary at $Q = 20$, beyond which the total energy captured by the ecosystem exceeds the available supply. As an observation here, partial communities become feasible when one component of $\boldsymbol{s}$ is disproportionately small.

### eferences

[1] Svirezhev YM, Logofet DO. Stability of Biological Communities. Mir Publishers; 1983.

[2] Hofbauer J, Sigmund K. Evolutionary Games and Population Dynamics. Cambridge University Press, Cambridge; 1998.

[3] Boyd SP, Vandenberghe L. Convex optimization. Cambridge university press; 2004.
